# Sex dependent glial-specific changes in the chromatin accessibility landscape in late-onset Alzheimer’s disease brains

**DOI:** 10.1101/2021.04.07.438835

**Authors:** Julio Barrera, Lingyun Song, Alexias Safi, Young Yun, Melanie E. Garrett, Julia Gamache, Ivana Premasinghe, Daniel Sprague, Danielle Chipman, Jeffrey Li, Hélène Fradin, Karen Soldano, Raluca Gordân, Allison E. Ashley-Koch, Gregory E. Crawford, Ornit Chiba-Falek

**Affiliations:** Division of Translational Brain Sciences, Department of Neurology, Duke University Medical Center, Durham, NC, 27710, USA; Center for Genomic and Computational Biology, Duke University Medical Center, Durham, NC, 27708, USA; Duke Molecular Physiology Institute, Duke University Medical Center, Durham, NC, 27701, USA; Department of Biostatistics and Bioinformatics, Duke University Medical Center, Durham, NC, 27705, USA; Department of Computer Science, Duke University, Durham, NC, 27705, USA; Department of Medicine, Duke University Medical Center, Durham, NC, 27708, USA; Department of Pediatrics, Division of Medical Genetics, Duke University Medical Center, Durham, NC, 27708; Center for Advanced Genomic Technologies, Duke University Medical Center, Durham, NC, 27708, USA

**Author notes:** These authors contributed equally to this work (Julio Barrera, Lingyun Song). To whom correspondence should be addressed: Ornit Chiba-Falek, Dept of Neurology, DUMC Box 2900, Duke University Medical Center, Durham, North Carolina 27710, USA, Gregory E. Crawford, Dept of Pediatrics, DUMC Box 3382, Duke University Medical Center, Durham, North Carolina 27708, USA., Allison E. Ashley-Koch, Dept of Medicine, DUMC Box 104775, Duke University Medical Center, Durham, North Carolina 27701, USA.

**Keywords:** ATAC-seq, chromatin accessibility, nuclei sorting, late-onset Alzheimer’s disease, snRNA-seq, gene dysregulation

## Abstract

In the post-GWAS era, there is an unmet need to decode the underpinning genetic etiologies of late-onset Alzheimer’s disease (LOAD) and translate the associations to causation. Toward that goal, we conducted ATAC-seq profiling using neuronal nuclear protein (NeuN) sorted-nuclei from 40 frozen brain tissues to determine LOAD-specific changes in chromatin accessibility landscape in a cell-type specific manner. We identified 211 LOAD-specific differential chromatin accessibility sites in neuronal-nuclei, four of which overlapped with LOAD-GWAS regions (±100kb of SNP). While the non-neuronal nuclei did not show LOAD-specific differences, stratification by sex identified 842 LOAD-specific chromatin accessibility sites in females. Seven of these sex-dependent sites in the non-neuronal samples overlapped LOAD-GWAS regions including *APOE*. LOAD loci were functionally validated using single-nuclei RNA-seq datasets. In conclusion, using brain sorted-nuclei enabled the identification of sex-dependent cell type-specific LOAD alterations in chromatin structure. These findings enhance the interpretation of LOAD-GWAS discoveries, provide potential pathomechanisms, and suggest novel LOAD-loci. Furthermore, our results convey mechanistic insights into sex differences in LOAD risk and clinicopathology.

**GRAPHICAL ABSTRACT:** 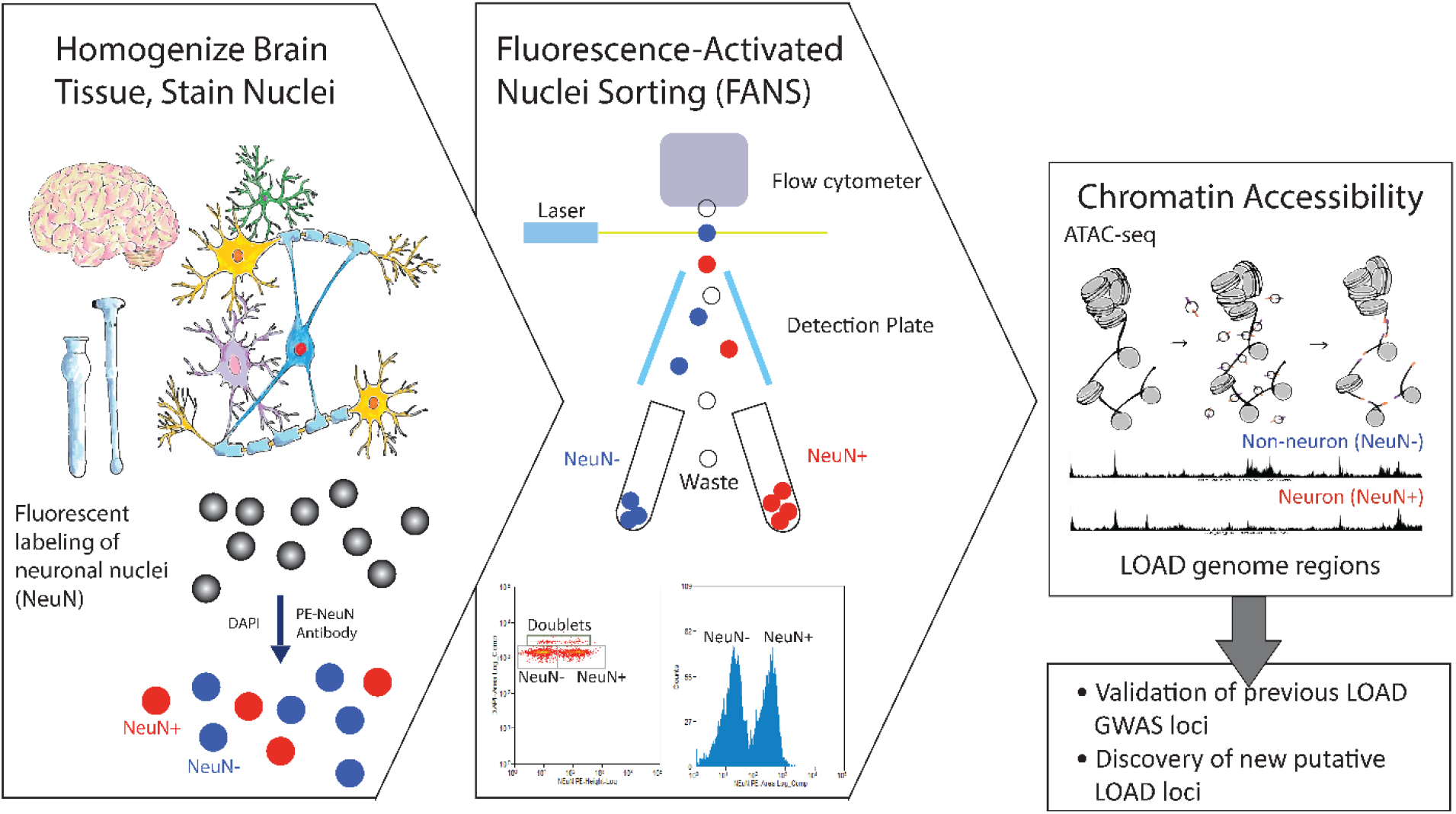

## Introduction

Large multi-center genome-wide association studies (GWAS) have identified associations between numerous genomic loci and late-onset Alzheimer’s disease (LOAD)^1–6^. The most recent GWAS meta-analysis reported a total of 25 LOAD-GWAS regions^7^. However, the precise disease-responsible genes, the specific causal genetic variants, and the molecular mechanisms of action that mediate their pathogenic effects are yet to be explained. Most LOAD-GWAS SNPs are in noncoding (intergenic and intragenic) genomic regions, and thus may have a gene regulatory function. Supporting this hypothesis, differential gene expression in brain tissues have been described between LOAD and healthy controls ^8,9, 10^. Moreover, several expression quantitative trait loci (eQTL) studies in brain tissues from cognitively normal^11^ and LOAD^12–15^ individuals reported overlaps with LOAD-GWAS loci. Collectively, these observations suggest that dysregulation of gene expression plays an important role in LOAD pathogenesis^16^. Noteworthy, integration of findings from LOAD epigenome wide association studies (EWAS) and GWAS also identified a number of shared loci^17–24^, providing further support for the role of gene dysregulation in LOAD pathogenesis.

To date, most brain expression, DNA-methylation, and chromatin studies have used brain tissue homogenates that represent multiple cell-types, *i.e*. various types of neurons and types of glia cells. The heterogeneity of brain tissue makes it difficult to determine the specific brain cell-type responsible for the changes in gene expression and in epigenome landscape. The mixture of cell-types could potentially mask signals corresponding to a particular cell-type, especially if the causal cell types comprise a small fraction of the entire sample. An additional shortcoming of studying bulk brain tissues is the bias related to sample-to-sample differences in the cell-type composition of the tissue. The obstacle of variability in the neuron:glia proportions across samples is even more pronounced when analyzing disease-affected brain tissues that underwent the neurodegeneration processes of neuronal loss and gliosis. Single cell experimental approaches can circumvent these limitations; however, these methods are costly for studying larger sample sizes and have been underutilized in the field of LOAD genetics. Frozen tissues pose additional technical challenges as it is difficult to isolate intact cell bodies. Fluorescence Activated Nuclei Sorting (FANS) was developed to extract, purify and sort nuclei (vs. cells) from frozen brain tissues^25^ using nuclei neuronal markers, such as NeuN, to greatly reduce cellular heterogeneity found in bulk tissues, and allow characterization of neuronal (NeuN+) and non-neuronal (NeuN-) cell populations. Recently, two studies used FANS to analyze LOAD-specific changes in DNA-methylation on a whole-genome level^26^ and across the *APOE* locus^27^. Furthermore, two new studies used single-cell (sc)RNA-seq from cortex of LOAD patients. The first found that the strongest LOAD-associated changes appeared early in pathological progression and were highly cell-type specific^28^, and the second identified LOAD-associated gene dysregulation in specific cell subpopulations, particularly for *APOE* and transcription factors^29^. These results further highlight the importance of cell-type specific analysis of human brain tissues to understanding of the molecular underpinnings and cellular basis of LOAD.

In this study, we performed NeuN-FANS from 40 archived frozen human brain samples (19 LOAD, 21 normal), and used the assay for transposase-accessible chromatin using sequencing (ATAC-seq) to characterize the chromatin accessibility landscape in neuronal and non-neuronal nuclei. We identified over 170,000 chromatin accessibility differences between neuronal and non-neuronal nuclei. We also report LOAD-specific differences in chromatin accessibility in both neurons and non-neurons. Interestingly, while the neuronal changes appeared to be independent of sex, in the non-neuronal cells LOAD differences in chromatin accessibility were detected only in females. LOAD-specific differences in chromatin accessibility significantly overlap with known LOAD GWAS regions, and also point to new candidate LOAD loci. These results provide new insights into the mechanistic and sex-specific pathogenesis of this disorder.

## Material and Methods

### Human Brain Tissue Samples

Following quality-control filtering, the final dataset was generated using fresh-frozen temporal cortex from neurologically healthy controls (n=21), mild LOAD (n=16), and severe LOAD (n=3) patients. These samples were obtained from the Kathleen Price Bryan Brain Bank (KPBBB) at Duke University, and the demographics for this cohort are included in **Table 1** and detailed in **Table S2**. Clinical diagnosis of LOAD was pathologically confirmed using Braak staging (AT8 immunostaining) and amyloid deposition assessment (4G8 immunostaining) for all LOAD samples. Braak staging was used to define mild (stages IV and below) vs. severe (stages V and VI) LOAD. All donors are Caucasians with APOE 33; post mortem interval (PMI) averaged 7.15 hours (standard error of the mean (SEM) 0.81). The archived frozen tissues have high-quality DNA as required for genomic analyses, and RNA (RIN≥8) suitable for quantitative RNA analyses. The project was approved by the Duke Institutional Review Board (IRB). The methods described were carried out in accordance with the relevant guidelines and regulations. Tissue samples were processed in random pairs of one normal and one LOAD patient. Tissue homogenization, nuclei extraction, FANS, and tagmentation were performed on each pair on the same day. Library preparation and sequencing was performed blinded to age, sex, and pathology. Out of this group of samples four female donors were analyzed by snRNA-seq, two of which were neurologically healthy controls and two age matched mild LOAD patients (Braak Stage III). These four samples were ages 79-90 with PMI averaged 8.02 hours (SEM=1.99) (**Table S2,** marked in *). Tissue samples were processed for snRNA-seq on the same day and in the same 10X Genomics microfluidics chip.

**Table 1:** Demographic description of study cohort. Abbreviations: mild Alzheimer’s disease (mAD), severe Alzheimer’s disease (sAD)

| | Pathology | Individuals | Males (%) | Age (y) ± SD | PMI (h) ± SD | NeuN+ (% ± SD) |
| --- | --- | --- | --- | --- | --- | --- |
| 40 Paired (neuronal and non-neuronal) | Normal | 21 | 11 (52.4) | 81.10 ± 9.70 | 7.91 ± 5.54 | 35.27 ± 13.26 |
| | mAD | 16 | 10 (62.5) | 79.75 ± 7.23 | 7.09 ± 5.58 | 38.59 ± 14.70 |
| | sAD | 3 | 1 (33.3) | 76.67 ± 7.77 | 5.43 ± 1.65 | 19.16 ± 7.49 |
| | Total | 40 | 22 (55) | 80.23 ± 8.49 | 7.41 ± 5.26 | 35.39 ± 14.16 |
| 9 Unpaired (non-neuronal only) | Normal | 3 | 0 | 84.67 ± 2.52 | 13.03 ± 4.95 | N/A |
| | mAD | 6 | 0 | 76.50 ± 8.48 | 16.95 ± 16.09 | N/A |
| | Total | 9 | 0 | 79.22 ± 8.73 | 15.64 ± 14.64 | N/A |

### Tissue dissociation and nuclei extraction

Methods were performed according to established protocols^25, 73, 74^ with some modifications. Briefly, 50 mg of frozen temporal cortex (gray matter) was thawed for 10 minutes on ice in lysis buffer (0.32M Sucrose, 5 mM CaCl2, 3 mM magnesium acetate, 0.1 mM EDTA, 10 mM Tris-HCl pH8, 1 mM DTT, 0.1% Triton X-100). The tissue was gently dissociated and homogenized in a 7 ml dounce tissue homogenizer (Corning) with approximately 25 strokes of pestle A in 45 seconds, then filtered through a 100 μm cell strainer. The filtered lysate was transferred to a 14 x 89 mm polypropylene ultracentrifuge tube, carefully underlaid with sucrose solution (1.8 M sucrose, 3 mM magnesium acetate, 1 mM DTT, 10 mM Tris-HCl, pH 8) and subjected to ultracentrifugation at 107,000 RCF for approximately 30 minutes at 4°C. Supernatant and the debris interphase were carefully aspirated, and 100 ul PBS (-Mg2+, -Ca2+) was added to the nuclei pellet. After a 5-minute incubation on ice, nuclei were gently resuspended and transferred to a 1.5 ml polypropylene microcentrifuge tube for staining.

For snRNA-seq downstream experiment, the above protocol was modified to optimize sample preparation for single-nuclei sorting. Briefly, 50mg frozen temporal cortex (gray matter) was thawed for 20 minutes on ice in lysis buffer and ultracentrifuged at 107,000 RCF for approximately 10 minutes at 4°C. Supernatant and the debris interphase were carefully aspirated, and 500 ul wash and resuspension buffer (1X PBS, 1% BSA, 0.2U/ul RNase Inhibitor) was added to the nuclei pellet. After a 5-minute incubation on ice, nuclei were gently resuspended and centrifuged at 300 RCF for 5 minutes at 4°C. The supernatant was again aspirated and 500 ul wash and resuspension buffer without RNase Inhibitor (1X PBS, 1% BSA) was added to the nuclei pellet. After a 1-minute incubation on ice, the nuclei were gently resuspended and filtered through a 35 um strainer. 10 ul nuclei were taken for quality assessment and counting prior to library preparation.

### Immunostaining of nuclei

Nuclei were stained in 0.05% BSA, 1% normal goat serum, DAPI (1μg/ml), and PE-conjugated anti-NeuN antibody (1:125, Millipore FCMAB317PE) in PBS (-Mg2+, -Ca2+), in the dark for 30 minutes at 4 °C. A DAPI-only control was prepared to set gates for sorting. After staining, nuclei were filtered through a 40 um cell strainer into a polypropylene round-bottom 5ml tube and sorted.

### Immunofluorescence microscopy

After homogenization and sucrose gradient ultracentrifugation, a portion of the nuclei was counted, resuspended in 4% PFA, stained, plated on 12 mm coverslips at 10,000 nuclei per coverslip, incubated 20 minutes at room temperature, mounted, and imaged on a confocal microscope.

### Fluorescence-activated nuclei sorting (FANS) of neuronal and non-neuronal nuclei

Sorting was performed using a MoFlo Astrios flow cytometer (Beckman Coulter) equipped with a 70μm nozzle, operating at 35 psi. Standard gating procedures were used. Briefly, the first gate allowed separation of intact nuclei from debris. The second gate allowed us to identify individual nuclei, and exclude doublets and other aggregates. The third and fourth gates distinguish between PE+ and PE-nuclei and allowed us to sort and separate NeuN+ nuclei from NeuN-nuclei. Nuclei were sorted into 1 ml PBS (-Mg2+, -Ca2+) in a 2 ml polypropylene tube pre-coated with 200 ul of 5% BSA and rotated at 20 rpm at 4C.

### Omni-ATAC on FANS nuclei

Approximately 100,000-700,000 sorted nuclei **(Table S2)** were used for ATAC-seq library preparation as described in the Omni-ATAC protocol^75^. Libraries were quantified by Qubit, and size distribution was inspected by Bioanalyzer (Agilent Genomic DNA chip, Agilent Technologies). Barcoded ATAC-seq libraries were combined into pools of 6 libraries and sequenced on an Illumina HiSeq 4000 sequencer (50 bp, single read) at the Duke Sequencing and Genomic Technologies shared resource.

### Omni-ATAC-seq on bulk tissue

In addition to performing ATAC-seq on NeuN+/- sorted nuclei, we also compared to ATAC-seq performed on total nuclei isolated from frozen tissue. Approximately 50,000 nuclei from 25 mg of pulverized frozen tissue were used for the transposition reaction applying the Omni-ATAC protocol as described previously ^75^.

### Data processing pipeline

ATAC-seq libraries made from 83 glia samples and 80 neuron samples were sequenced on Hi-seq 4000. Raw fastq sequencing files were first processed through cutadapt (v 1.9.1) to remove adaptors and bases with quality scores < 30. Filtered reads were aligned to hg19 by Bowtie2 (v 2.1.0) using default parameters^76^. Bam files were sorted by samtools (v 0.1.18) and duplicates were removed by Picard MarkDuplicates. Sequences that overlapped ENCODE “blacklist” regions (https://sites.google.com/site/anshulkundaje/projects/blacklists) were removed, and narrow open chromatin peaks were called by Model-based Analysis of ChIP-seq (MACS v 2.1), with parameters --nomodel --shift −100 --ext 200, using a FDR cutoff of q < 0.01 or 0.05^77^. For visualization on the UCSC browser, BigWig files were generated using wigToBigWig (v 4).

### Data quality control

We characterized the quality of all ATAC-seq datasets by the following metrics, based on ENCODE suggestions^78^ https://genome.ucsc.edu/ENCODE/analysis.html **(Table S1)**: (1) total numbers of reads, (2) numbers of reads trimmed by cutadapt, (3) total bases entering cutadapt, (4) quality-trimmed bases by cutadapt, (5) total numbers of reads processed by Bowtie2, (6) uniquely mappable reads, (7) percentages of alignment, (8) read aligned to the mitochondrial chromosome, (9) peak calls at FDR q < 0.05, (10) peak calls normalized by sequencing depth, (11) GC content of sequences, (12) Non-Redundant Fraction (NRF), which is equal to the number of distinct uniquely mapped reads divided by total reads, (13) PCR Bottleneck Coefficient 1 (PBC1), and (14) PCR Bottleneck Coefficient 2 (PBC2). To reduce the impact of samples with lower quality, we removed datasets with normalized peak calls at FDR q < 0.01 < 100, and any additional replicates that displayed lower signal-to-noise ratios. The 90 remaining datasets were analyzed by PCA and hierarchical clustering (hclust function, method=“ward.D”), of which one NeuN-sample was identified as an outlier **(Fig. S10)**. After excluding this outlier, we obtained a final set of 89 datasets from 49 donors (40 donors with matching NeuN+ and NeuN-data, and 9 donors that only contained NeuN-data). Sequencing and QC metrics for all 89 samples are described in **Table S2**. The library complexity for these 89 samples are comparable to ENCODE Data Quality Metrics spreadsheet published on 2012-04-25 (https://genome.ucsc.edu/ENCODE)

### Evaluation of variables affecting chromatin peaks

Prior to the differential analyses described below, we considered the effect of 37 variables that could impact the quality of the ATAC-seq results for each sample. These variables were collected from key features of the ATAC-seq processing pipeline, as well as individual sample characteristics such as case-control status, sex, and age. For example, the metadata for each sample included transposase batch, nuclei sorting date, mass, mean GC percentage of sequenced reads, mean mapped read length, alignment quality metrics, subject age at death, sex, diagnosis, and postmortem interval (PMI). Two subjects were missing PMI, therefore we used the R package MICE^79^ to impute missing values using the classification and regression trees methodology.

### Covariate selection

For differential peak analyses, selection of covariates for adjustment was carefully tailored to each comparative analysis to account for the variable number of peaks between groups and to minimize false positives in peak calling. We performed a linear regression of all metadata variables against the first 10 principal components (PCs) of the TMM normalized peak quantification in an effort to identify covariates to be utilized in the differential expression analyses. In an iterative process, we selected one variable (preferentially a variable directly related to the ATAC-seq experiment, explaining one of the largest proportions of variance and with few parameters), regressed its effect on the peak quantifications and performed a new principal component analysis independent of the selected variable(s). We repeated this procedure until there were no more Bonferroni significant (q<0.05) variables associated with the peak PCs. For the comparison of neuronal and non-neuronal nuclei, we selected the following variables for our differential chromatin analysis: NRF, number of nuclei, normalized peak calls, nuclei sort date, alignment percentage, passing base pairs, age, and donor. For the comparison of LOAD vs. control neuronal nuclei, we selected normalized peak calls, alignment percentage, nuclei sort date, and number of nuclei as covariates. For the comparison of LOAD vs. control non-neuronal nuclei, we included the number of nuclei, NRF, nuclei sort date, and alignment percentage as covariates in the differential chromatin analysis. When performing a sex-stratified analysis of LOAD vs. control samples in the NeuN+ nuclei, we selected alignment percentage in the female-only subset and nuclei sort date in the male-only subset. Finally, in the sex-stratified differential chromatin analysis of LOAD vs. control NeuN-nuclei, we included normalized peak calls as a covariate in the female-only subset, and normalized peak calls and percent GC content as covariates in the male-only subset. In the supplemented comparison of LOAD vs. control NeuN-nuclei in females only (nine additional non-neuronal samples), percent GC content was included as a covariate.

### Differential chromatin accessibility analyses

Differential ATAC-seq peaks were detected using EdgeR package (version 3.22.3), which models counts using a negative binomial distribution^80^. ATAC-seq reads corresponding to chrX and chrY were excluded due to unequal numbers of female and male donors. For each comparison, counts of reads within peaks which were merged from all of the replicates were extracted from BigWig files and normalized by weighted trimmed mean of M-values^81^ (TMM). Quasi-likelihood F-tests (QLF) was performed to determine differential sites at cut off adjusted p values < 0.05. As an approximate error model, QLF works more robustly and gives more reliable Type I error rate control than the other options, especially when there are smaller numbers of replicates (EdgeR User Guides, Bioconductor package vignettes). Three levels of case-to-control chromatin accessibility comparisons were performed **(Fig. 1g)**: ***Level_1***: 40 NeuN-vs 40 NeuN+, all samples are from matched donors; ***Level_2***: (A) NeuN-, 19 cases vs 21 controls (B) NeuN+, 19 cases vs 21 controls; ***Level_3***: (A) NeuN-females, 11 cases vs 11 controls; (B) NeuN-males 8 cases vs 10 controls; (C) NeuN+ females, 11 cases vs 11 controls; and (D) NeuN+ males, 8 cases vs 10 controls. Since the number of female samples was less than that of male, we included 9 additional NeuN-datasets (those that did not have matching NeuN+ data from the same donor but met QC criteria) into Level 3 and processed an additional comparison: (E) NeuN-females,14 cases vs 13 control. For all these comparisons we included covariates (described in “covariate selection”) in the EdgeR analysis with model design as open chromatin accessibility ~ Disease (LOAD state) + covariates. Importantly, for each analysis, we remerged and recalled the chromatin peaks. Thus, the separate analyses were not direct subsets of each other.

**Fig. 1:**
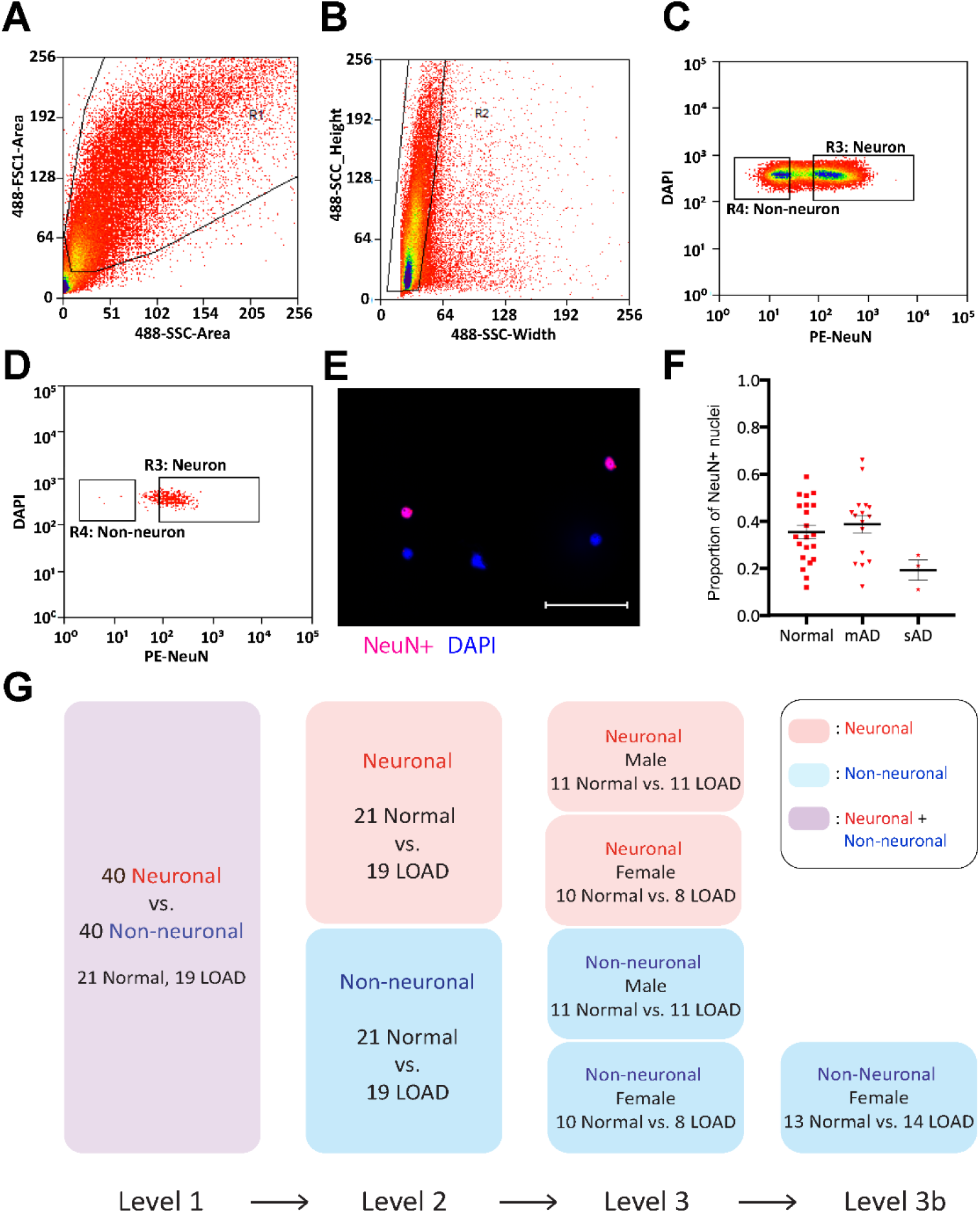
Isolation of nuclei from frozen brain samples and analysis of ATAC-seq data. Human postmortem frontal cortex was dissociated, nuclei were isolated and stained with the nuclear stain DAPI and a monoclonal NeuN antibody conjugated to PE. **(A)** Nuclei were first sorted based on their forward and side scatter from all possible events (R1 gate). **(B)** Single nuclei were further sorted based on their size from the doublets or larger clumps of nuclei (R2 gate). **(C)** DAPI positive single cells were gated as either NeuN-PE positive (neurons, R3 gate) or NeuN-PE negative (glia, R4 gate). **(D)** Post-sort data showing the purity of the separation between neuronal and non-neuronal nuclei. **(E)** Fluorescence image showing unsorted nuclei stained for NeuN (red) and DAPI (blue). The scale bar represents 100um. **(F)** Proportion of neuronal nuclei from each sample. Error bars show standard error of the mean. **(G)** Overview schematic of Levels 1, 2, 3, and 3b of differential analysis. Level 1 compares neuronal vs. non-neuronal for 21 normal and 19 LOAD samples. Level 2 compares normal vs. LOAD for each neuronal and non-neuronal subpopulation. Level 3 compares LOAD samples separated by female and male. Level 3b is the same comparison done after adding 9 female non-neuronal samples (3 normal and 6 LOAD).

### Genomic distribution analysis

ATAC-seq peaks were categorized as promoters (transcription start site to upstream 1kb), 1^st^ exon, intragenic (excluding 1^st^ exon), 5’ UTR, 3’ UTR, and intergenic, based on human gene hg19 NCBI RefSeq gene information from the UCSC genome browser.

### Gene ontology analysis

Differential open chromatin regions were characterized by GREAT (v 3.0.0) (http://bejerano.stanford.edu/great/public/html/), using the hg19 genome as background regions. Genes associated to open chromatin regions were determined by default “basal plus extension” settings (i.e., 5 kb upstream, 1 kb downstream, plus distal up to 1000 kb).

### Overlap with LOAD GWAS loci

The tag SNP for 25 LOAD loci were obtained from the literature^7^. Genome coordinates +/100kb surrounding each GWAS tag SNP were used to identify any differential open chromatin region that mapped within the region. Permutation analyses were performed by randomly selecting the same numbers of sites from the union set of ATAC-seq peak calls. For each comparison, we performed 10,000 permutations to estimate the empirical P-value.

### Motif search

Transcription factor motif enrichments were evaluated using HOMER^82^ (v4.10.3) and MEME Suite^83^. For each comparison, we used the centered 300 bp of open chromatin regions as input, and the union peak calls as background. GC matching was applied to the background peaks to ensure that this did not lead to spurious results. We primarily used all peaks as the background, but to check for accuracy, we also randomly selected 10,000 peaks as the background from all peaks. For this analysis, we focused on the known motif search rather than search for *de novo* motifs.

### Differential expression analysis of ROSMAP data

Gene expression data from AMP-AD was obtained from the Religious Orders Study and Memory and Aging Project (ROSMAP)^84, 85^. This study includes RNA-seq data from the dorsolateral prefrontal cortex of 724 subjects^86, 87^ limma was used to perform the differential analysis in R on normalized FPKM values obtained from RNA-seq of ROSMAP bulk tissue samples. Samples with cogdx of 1 (no cognitive impairment, n=201) and 4 (Alzheimer’s dementia and no other case of cognitive impairment, n=222) were included in the analysis.

### Single-nuclei library preparation and sequencing

Libraries were prepared for snRNA-seq using the Chromium Single Cell 3’ Reagent Kits v3 (10X Genomics) according to the manufacturer’s instructions. Nuclei were diluted in nuclease-free water to a concentration of 1 million nuclei / ml in a final volume of 100 ul, and transferred to an 8-strip tube on ice. 16,000 nuclei were added to the Master Mix for a targeted nuclei recovery of 10,000. Libraries were pooled onto a single S1 flow cell, and sequenced using the Illumina NovaSeq 6000 system to obtain paired-end 2 x 100bp reads. Sequencing saturations ranged from 59.3% to 81.2%.

### snRNA-seq data analysis

*CellRanger* software version 3.1.0 (10X Genomics) was used to demultiplex raw Illumina base call files into FASTQ files. A pre-mRNA GTF file was generated with the pre-built GRCh38 3.0.0 human reference using the Linux utility *awk*, in order to be compatible with CellRanger and to include intronic reads from nuclear RNA in UMI counts for each gene and barcode. The CellRanger count pipeline was run to align reads to the pre-mRNA GRCh38 reference and gene expression matrices generated separately for each of the four samples were merged together into a single matrix.

For quality control and filtering, the *quickPerCellQC* function from the R package *scater* v1.14.6^88^ was used to identify low-quality cells based on QC metrics (UMI count, number of genes detected, percentage of UMIs that mapped to the mitochondrial genome). Cells in which 229 or more genes were detected and less than 17.4% of UMIs mapped to the mitochondrial genome were used in downstream analyses. Out of 21,262 nuclei in the initial dataset 18,032 remained after filtering with a median number of 2,332 detected genes per nucleus. The R package Seurat v3.1.1 standard workflow was used for integration of multiple samples to combine the four samples into a unique dataset^89^. Prior to integration, gene expression was normalized for each sample by scaling by the total number of transcripts, multiplying by 10,000, and then log transforming (log-normalization). We then identified the 2,000 genes that were most variable across each sample, controlling for the relationship between mean expression and variance. Next, we identified anchor genes between pairs of samples using the *FindIntegrationAnchors* function that were then passed to the *IntegrateData* function to harmonize the four samples.

We scaled the integrated dataset before running a Principal Component Analysis (PCA). To distinguish principal components (PCs) for further analysis, we used the *JackStraw* method to determine statistically significant PCs and found that up to 30 PCs were enriched in genes with a PC score that was unlikely to have been observed by chance. We then utilized the shared nearest neighbor (SNN) modularity optimization-based clustering algorithm implemented in Seurat for identifying clusters of cells. This was performed using the *FindNeighbors* function with 30 PCs, followed by the *FindClusters* function with the Louvain algorithm using a 0.4 resolution. This allowed us to assign cells into a total of 21 clusters. We applied the uniform manifold approximation and projection (UMAP) method on the cell loadings of the previously selected 30 PCs to visualize the cells in two dimensions and to separate nuclei into clusters.

Differential expression to identify cluster markers that are conserved between the samples was performed using the Seurat function *FindConservedMarkers* for each cluster on the normalized gene expression before integration. R package SingleR v1.0.1^90^ was used to annotate cell types based on correlation profiles with 713 bulk microarray samples from the Human Primary Cell Atlas^91^ (HPCA) as reference expression data. Four major cell types were detected by the SingleR method using HPCA: macrophages, astrocytes, neurons, and endothelial cells. Because of the specific expression of microglial markers PTPRC and CSF1R in the macrophage cluster, as well as the differences in biological systems used in the HPCA reference, we manually refined the HPCA annotation of this specific cluster to microglia. Identification of neuronal vs. non-neuronal clusters for some differential expression analyses was performed by determining the proportion of cells expressing the neuronal marker NeuN/RBFOX3. Clusters with more than 50% of cells expressing NeuN/RBFOX3 were defined as neuronal clusters. Control samples showed an average of 2,417 (SD 169) neuronal and 2,319 (SD 1,149) non-neuronal nuclei, whereas LOAD samples showed 1,329 (SD 294) neuronal and 2,952 (SD 1,783) non-neuronal nuclei. NeuN neuronal clusters and neuronal clusters identified by SingleR using the HPCA reference were very consistent with only one additional cluster in NeuN neuronal clusters (out of a total of 16 clusters). Differential expression analyses between female LOAD nuclei and control nuclei within specific groups of nuclei were performed using the Wilcoxon Rank Sum test as implemented in the *FindMarkers* function in the Seurat package. ‘min.pct’ and ‘logfc threshold’ arguments were set to 0 to allow for the testing of a majority of genes in each analysis.

## Results

### Study sample and quality control analyses

Nuclei isolated from archived frozen temporal cortex of 51 individuals with LOAD (n=26) or controls (n=25) **(Table S1)** were stained and sorted using a PE-conjugated monoclonal NeuN antibody **(Fig. 1a-d)**. Staining of pre-sorted nuclei with nuclear membrane markers was confirmed by immunofluorescence **(Fig. 1e)**. We observed a smaller proportion of sorted neuronal (NeuN+) nuclei from total isolated nuclei in the severe LOAD cases **(Table 1, Fig. 1f)**, as expected due to neuronal cell loss and gliosis, hallmarks of LOAD pathological progression^30, 31^. We next performed ATAC-seq using the NeuN+ and NeuN-sorted nuclei populations. A total of 90 neuronal and non-neuronal datasets passed quality control criteria (see Methods), which were derived from 40 individuals (19 LOAD cases and 21 controls) that had matched neuronal and non-neuronal data, and an additional 9 individuals (3 LOAD cases and 6 controls) that had only non-neuronal data. The final cohort of 40 consisted of 22 males and 18 females with similar post-mortem intervals (PMI); all donors were Caucasians and homozygous for *APOE* e3 **(Table S2)**. Subsequent analyses compared different sub-groups **(Fig. 1g**). Correlations of all potential numerical covariates showed expected patterns of co-linearity (**Fig. S1**). The samples displayed a range of QC metrics, with some samples displaying a higher signal to noise (**Table S1)**. Repeated experiments on a subset of samples demonstrated that signal to noise was reproducible and thus a sample-specific characteristic (**Table S1**). In addition to randomizing sample preparation (see Methods), we found that no metadata variables were significantly associated with case-control status (**Fig. S2**), indicating an absence of batch effects.

### Differential analysis of ATAC-seq data between neuronal and non-neuronal nuclei

To demonstrate that we can robustly identify chromatin accessibility differences between major cell types of the brain, we first compared the ATAC-seq profiles between neuronal and non-neuronal nuclei groups from the entire study sample (Level 1, **Fig. 1g**). PCA of all samples showed that 37.5% of the total variance was explained by the cell type (neuronal versus non-neuronal, **Fig. S3, Fig. S11)**. Performing quantitative differential chromatin accessibility analysis using EdgeR (FDR q < 0.05), we found that 87,570 regions were more accessible in the neuronal population, 83,171 regions were more accessible in the non-neuronal population, and 54,484 regions were not detected as differential **(Fig. 2a, Table S10)**. Representative screenshots show differential ATAC-seq peaks around genes known to be expressed specifically in neuronal **(Fig. 2b)** or non-neuronal **(Fig. 2c)** cell types. A higher percentage of sites more accessible in non-neuronal population mapped to promoters, which could be explained by the more distal regulation of neuronal cell types or the greater diversity and heterogeneity of the cell types composing the non-neuronal population **(Fig. S4)**. Using GREAT gene ontology analysis ^32^, neuronal-specific regions were enriched with genes associated with neuronal function, while non-neuronal specific regions were enriched with genes implicated in glial function **(Table S3)**.

**Fig. 2:**
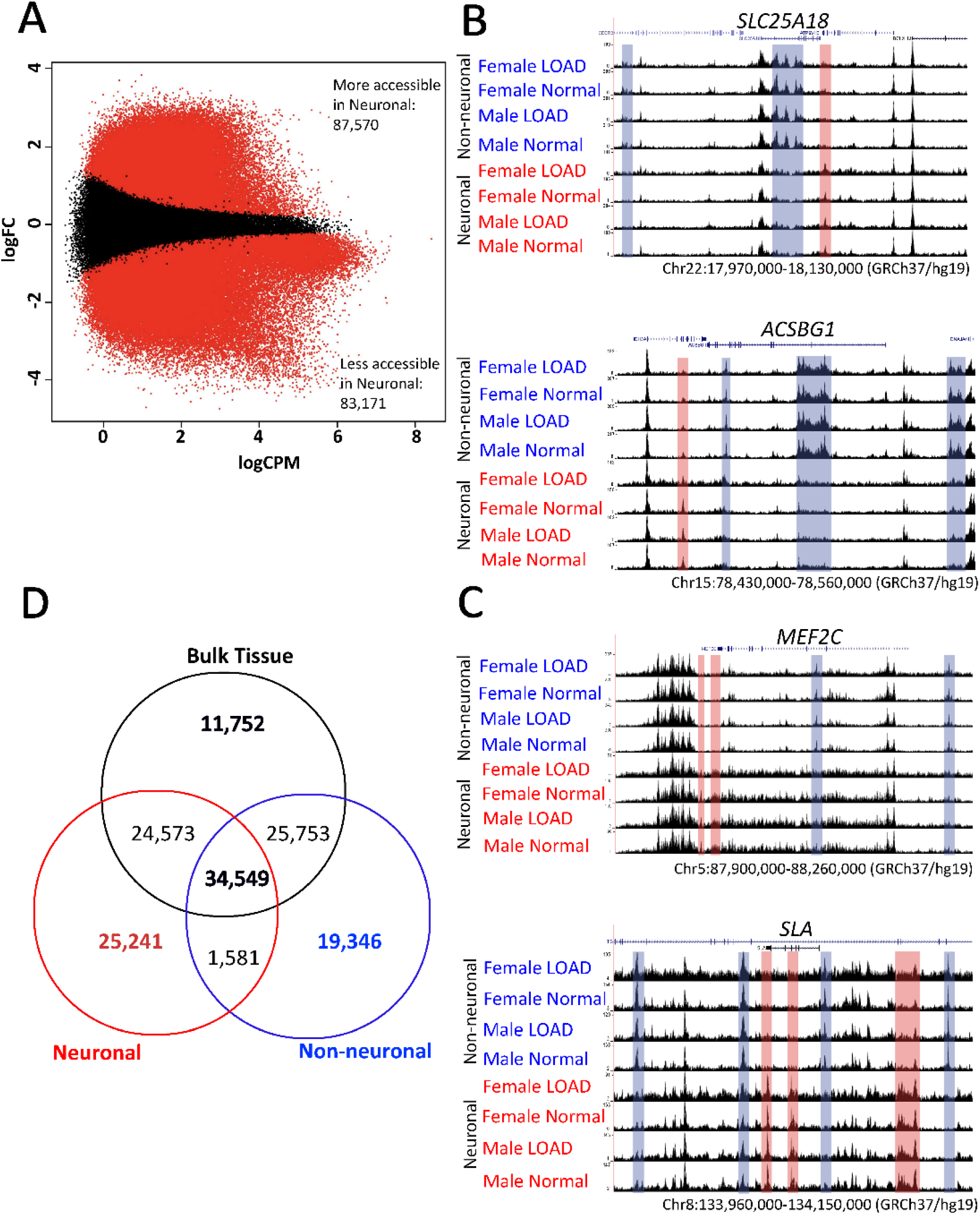
Level 1 comparison of ATAC-seq data from neuronal vs non-neuronal nuclei. **(a)** MA plot showing differential ATAC-seq sites between neuronal (blue) vs. non-neuronal regions (red). Red dots represent ATAC-seq peaks that are significantly different between groups (FDR q < 0.05). **(B)** ATAC-seq data around non-neuronal-specific genes *SLC25A18* (upper panel) and *ACSBG1* (lower panel). Boxes highlight peaks that are more accessible in neuronal (red) or non-neuronal (blue) nuclei. **(C)** ATAC-seq data around neuron-specific genes *MEF2C* (upper panel) and *SLA*(lower panel). All regions indicate hg19 coordinates. **(D)** Venn diagram of ATAC-seq peaks detected in whole tissue, sorted neuron and sorted non-neuron nuclei from 6 donor-matching samples.

Motif analysis for regions with increased chromatin accessibility in neuronal nuclei showed enrichment for the transcription factor recognition sites of Early Growth Response 2 (EGR2), Regulatory Factor X1 (RFX1), and Myocyte Enhancer Factor 2C (MEF2C)^33–35^, while motif analysis for regions with increased chromatin accessibility in non-neuronal nuclei showed enrichment for the SRY-related HMG-box (SOX) transcription factor family^36^ **(Table S4)**. When compared to a previous NeuN sorted-nuclei ATAC-seq study^37^ using a smaller number (n = 8) of healthy brain samples, we found a substantial degree of overlap **(Fig. S5a)** with similar peak length distribution **(Fig. S5b)**. However, the 5x larger number of samples used in our study identified >80,000 additional significantly differential chromatin sites.

### Comparison of sorted nuclei vs. bulk brain tissue homogenate

To our knowledge, no study has directly compared chromatin accessibility differences between bulk brain tissue, neuronal, and non-neuronal fractions. Bulk brain ATAC-seq data was generated using pulverized frozen whole tissue homogenate samples from six individuals for whom high-quality ATAC-seq data was collected from neuronal and non-neuronal populations **(Table S5)**. Binary peak overlap analysis shows that while there are some regions that are only accessible in bulk tissue, a substantially larger number of sites are uniquely detected in the nuclei populations of neuronal and non-neuronal only **(Fig. 2d)**. This observation justifies the performance of ATAC-seq analysis by brain cell-type to reduce cellular heterogeneity as it allows the identification of more cell type specific signals. Quantitative differential chromatin accessibility (EdgeR comparisons **(Fig. S6)** and GO annotations **(Table S6-8)** showed that chromatin accessibility sites specifically identified in bulk tissue, neuronal and non-neuronal populations displayed many biologically relevant pathways.

### Identification of LOAD-associated differences in chromatin accessibility

To determine the association of LOAD status with changes in chromatin accessibility, we performed differential ATAC-seq analysis of 19 LOAD compared to 21 healthy controls stratified by brain cell-type (neuronal and non-neuronal only populations; Level 2, **Fig. 1g)**. The comparison using the neuronal nuclei data resulted in 211 neuronal chromatin differences (FDR q < 0.05) between LOAD and control **(Fig. 3a, Table S11)**, while the analysis of the non-neuronal nuclei detected no differential chromatin between LOAD and control (**Fig. 3b**). Motif analysis using the 141 sites that showed decreased chromatin accessibility in LOAD neuronal nuclei were enriched for transcription factor motifs including Wilms Tumor protein (WT1), Early Growth Response 1 (Egr1), Retinoic acid receptor gamma (RARg), and Kruppel like factor 14 (KLF14) **(Fig. 3c)** that have been reported relevant to brain function^38–41^. No significant motif enrichment was detected from 70 sites (FDR q < 0.05) with increased chromatin accessibility in neuronal LOAD samples.

**Fig. 3:**
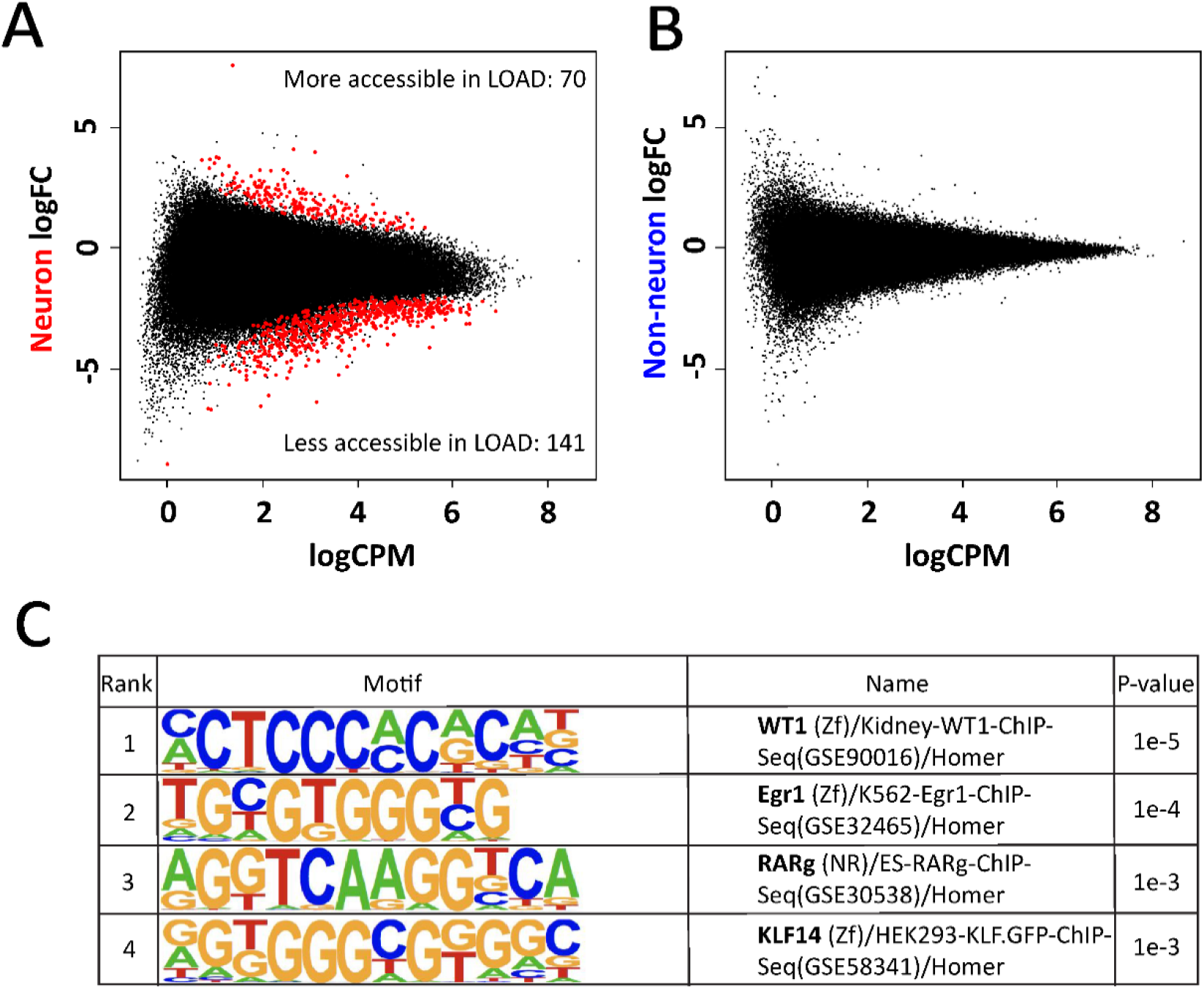
Level 2 comparison of ATAC-seq data between LOAD cases and controls. MA plots of differential chromatin sites for (A) neuronal and (B) non-neuronal nuclei. Red dots represent differential ATAC-seq sites with FDR q < 0.05. (C) Motifs that are enriched in neuronal ATAC-seq sites that are less accessible in LOAD samples. Size of red dots were increased for visibility.

### Identification of sex-dependent chromatin accessibility differences between LOAD and control samples

Since sex has an effect on LOAD onset and progression^42^, we performed a sex-stratified differential analysis of LOAD compared to control (Level 3, **Fig. 1g)** to examine whether the effect of sex on LOAD risk is mediated, at least in part, by chromatin remodeling. The differential analysis of the neuronal groups did not yield significant differences between LOAD and normal when stratified by sex for either females or males (FDR q > 0.05) **(Fig. 4a-b)**. Analysis of the non-neuronal nuclei identified 24 chromatin accessibility differences in female LOAD compared to control **(Fig. 4c, Table S12)**, while no significant differences were identified when the analysis was performed in males **(Fig. 4d)**. To further investigate this trend, we increased the sample size with 9 additional non-neuronal samples (6 female controls and 3 female LOAD cases). Differential analysis using ATAC-seq data from the larger female non-neuronal dataset resulted in 842 differential sites between LOAD and control (FDR q < 0.05) **(Fig. 4e, Table S13)**.

**Fig. 4:**
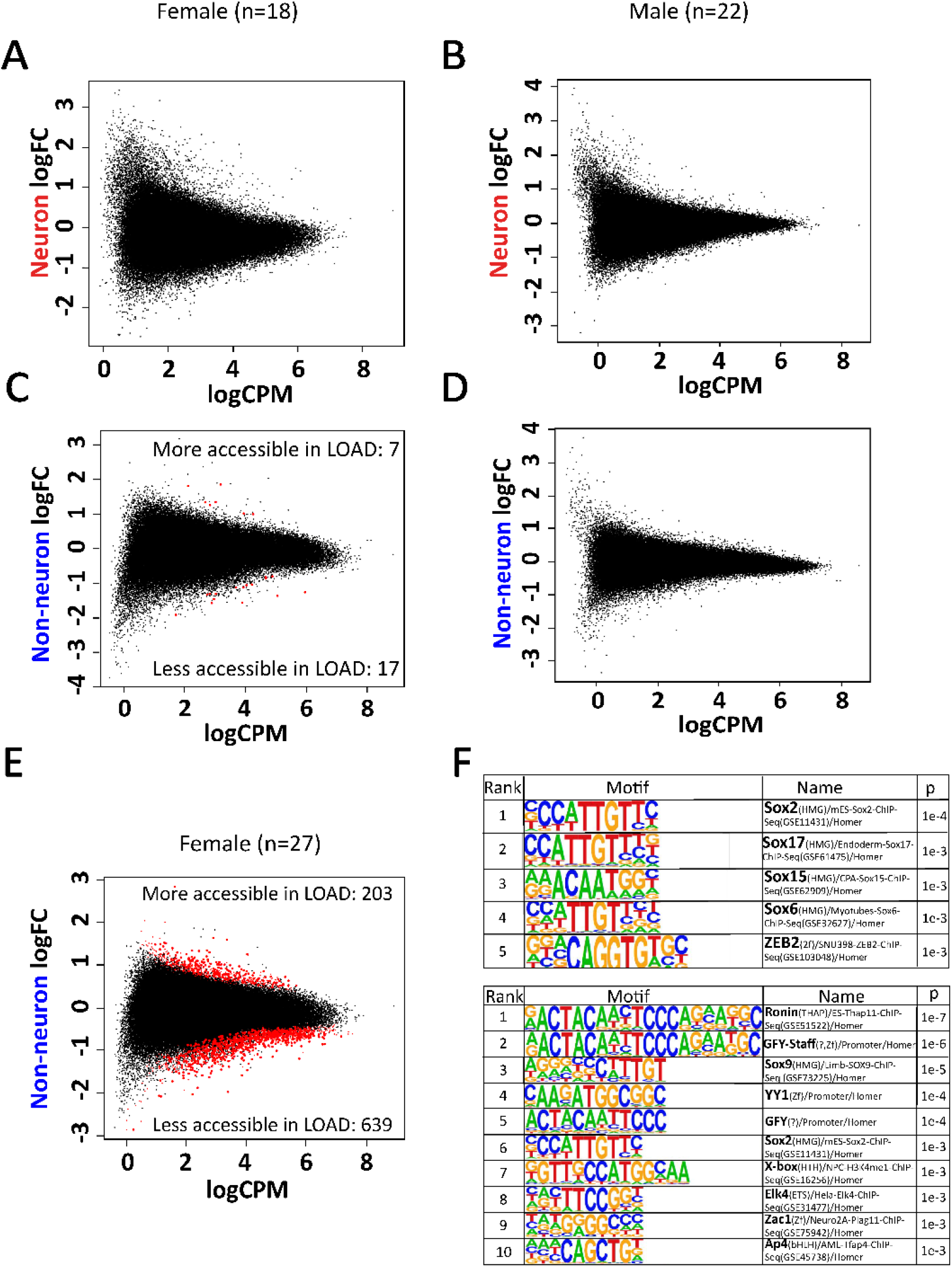
Level 3 comparison of ATAC-seq data between LOAD cases and controls separated by sex. MA plot of differential sites for **(A)** female neuron (n=18; FDR q < 0.05), **(B)** male neuron (n=22; FDR q < 0.05), **(C)** female non-neuron (n=18, FDR q < 0.05), **(D)** male non-neuron (n=22; FDR q < 0.05), and **(E)** female non-neuron with additional samples (n=27; FDR q < 0.05). **(F)** Motifs enriched for 203 sites that are more accessible in female non-neuronal LOAD and for 639 sites that are less accessible in female non-neuronal LOAD (FDR q < 0.05). Size of red dots were increased for visibility.

Comparison of these results with the LOAD associated chromatin accessibility sites obtained from the differential analysis of female neuronal nuclei showed that while there are differences detected in both female neuronal and glial cells in LOAD cases versus controls, we detected larger effect sizes in the female glia cells (**Fig. S12a**). In addition, several differential regions identified in female non-neuronal nuclei were also observed in male non-neuronal nuclei with similar trends, however, these did not reach statistical significance (**Fig. S12b**). Nonetheless, these associations in the female group showed a stronger effect size (**Fig. S12b**).

We performed motif enrichment using the 203 sites that were more accessible in non-neuronal female LOAD and found significantly enriched motifs for the SOX family **(Fig. 4f)**. The analysis of 639 sites that displayed decreased accessibility in non-neuronal female LOAD was enriched for transcription factors known to be highly associated with glia or neuron functions, such as RONIN, SOX9, YY1 and ELK4^43–46^ **(Fig. 4f)**.

To determine whether binding of transcription factors identified in female non-neuronal cells (Level 3, **Fig. 1g**) has downstream effects on expression of target genes, we used the publicly available ROSMAP bulk RNA-seq data to perform differential expression analyses of those genes for LOAD vs. Normal. We found that several genes exhibited changes in the expected direction, including *ZAC1* (*PLAGL1*), *YY1*, and *SOX2* (**Table S15**). This provides further mechanistic insights into gene dysregulation underlying LOAD pathogenesis in females. Last, we also used the chromatin accessibility differential sites identified in female non-neuronal LOAD (Level 3) for GO analysis and found pathways involved in immune response and myelination **(Table S9)**.

### Overlap of LOAD-specific differential chromatin accessibility sites with LOAD GWAS regions

To determine the relationship of the LOAD-specific chromatin accessibility sites we compared these data with LOAD GWAS regions^7^. We defined LOAD GWAS regions by anchoring on the top 25 associated SNPs +/- 100kb, and cataloged LOAD-GWAS regions of 200kb each. Importantly, we were comparing genomic regions of chromatin accessibility to the GWAS regions, not chromatin QTLs, to GWAS loci. Of the 211 LOAD-specific differential chromatin accessibility sites in neuronal samples (FDR q < 0.05, **Fig. 3a)**, we identified five sites that overlap four of the 25 LOAD GWAS regions **(Table 2)**. We show representative examples of overlapping LOAD-specific differential chromatin accessibility sites surrounding the *PTK2B* and *CLU* GWAS loci **(Fig. 5a, Fig. S7)**. Using permutation testing, we did not detect a significant enrichment with LOAD GWAS regions compared to regions selected at random **(Fig. S8a-b).**

**Table 2.**
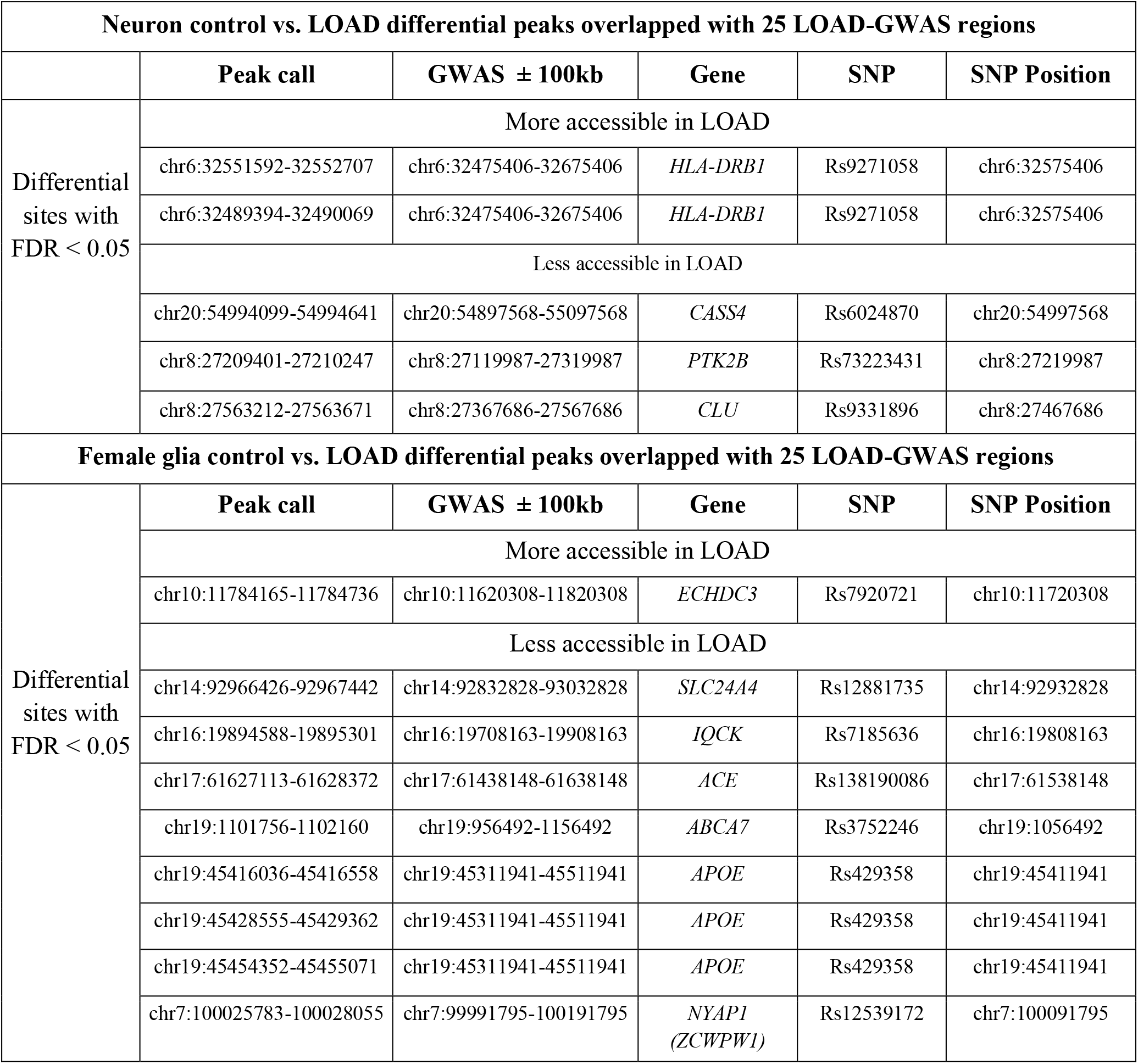
Neuron control vs LOAD differential peaks overlapped with 25 LOAD-GWAS regions as well as female glia control vs LOAD differential peaks overlapped with 25 LOAD-GWAS regions. For each panel, upper: more open in LOAD; lower: less open in LOAD

**Fig. 5:**
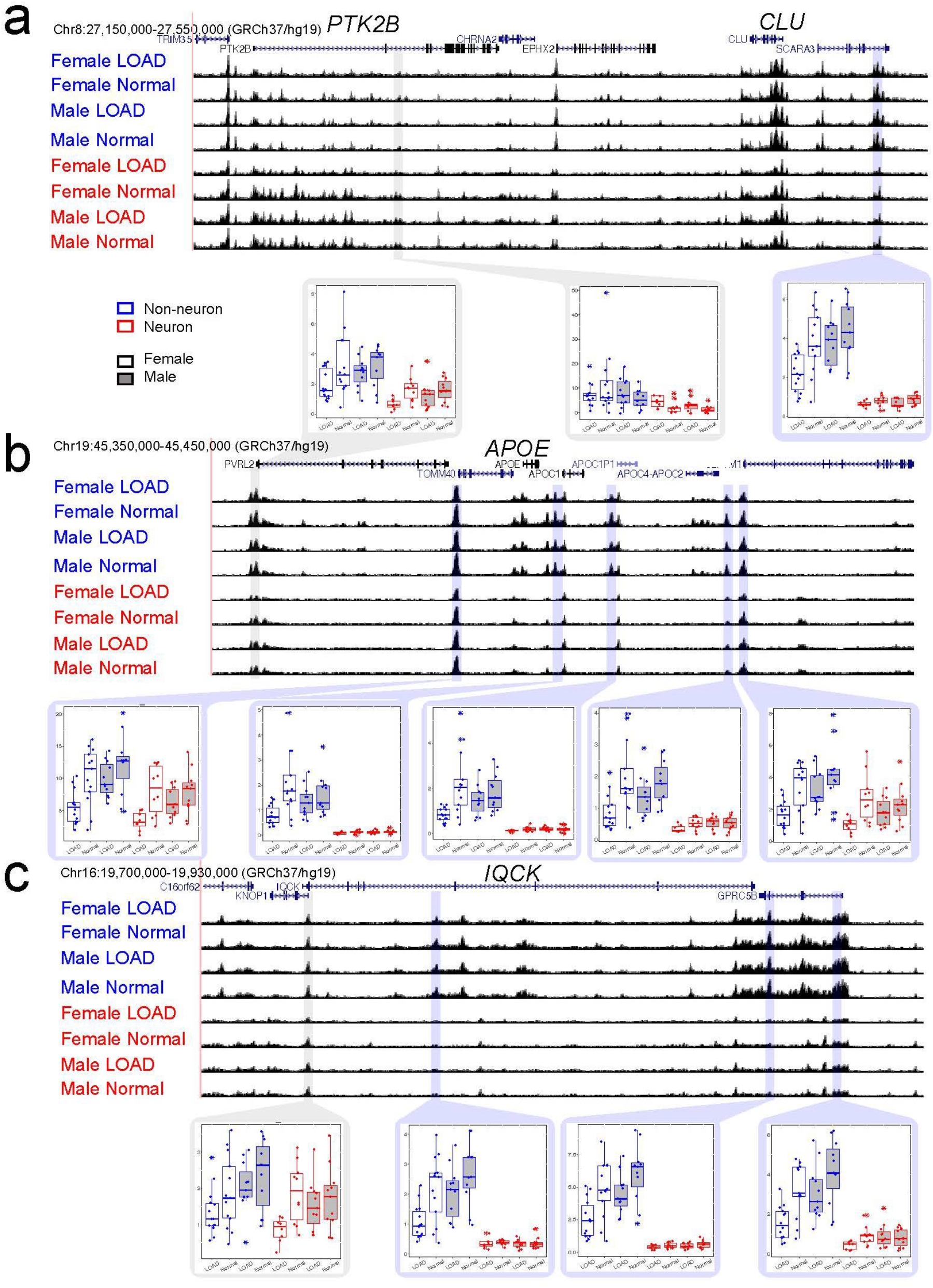
Differential LOAD specific ATAC-seq peaks around LOAD-GWAS regions. Screenshots of ATAC-seq data around **(A)** *PTK2B, CLU*, **(B)** *APOE*, and **(C)** *IQCK* loci. Box plots show ATAC-seq read counts for individual ATAC-seq peaks (blue frames highlight significant differential peaks for cases vs. controls, gray frames show control peaks that are not differential between cases and controls). Box plots are color coded for non-neuronal (blue) neuronal (red), female (no fill), and male (gray fill). All tracks show hg19 coordinates and all y-axes on tracks range from 0 to 250. For box plots, the line within each box represents the median, and the top and bottom borders of the box represent the 25 ^th^ and 75^th^ percentiles, respectively. The top and bottom whiskers of the box plots represent the 75^th^ percentile plus 1.5 times the interquartile range and the 25^th^ percentile minus 1.5 times the interquartile range, respectively.

Of the 842 differential sites in non-neuronal female LOAD (FDR q < 0.05, **Fig. 4e)**, we detected nine differential sites that overlap seven of the 25 LOAD-GWAS regions **(Table 2)**. Using permutation analysis (see Methods), we found significant enrichment in the amount of overlap with sites less accessible in non-neuronal female LOAD **(P < 0.05, Fig. S8c-d)**. Representative examples of differential chromatin accessibility sites between LOAD cases and controls overlapping representative LOAD-GWAS regions are shown for the loci *APOE* **(Fig. 5b, Fig. S7b)**, and *IQCK* **(Fig. 5c, Fig. S7c).** While differences around the *APOE* and *IQCK* locus are more pronounced in females, we also detect subtle differences in the same direction in males **(Fig. 5b-c, Fig. S7b-c).** The identification of LOAD differences in chromatin accessibility in the *APOE* region, despite all cases and controls being of *APOE* e3/e3 genotype, underscores our identification of a regulatory mechanism, rather than genetic mechanism, at this locus.

### Overlap of LOAD-specific differential chromatin accessibility sites in female non-neurons with single nuclei RNA-seq data

Next, we performed functional validation of key findings from the chromatin accessibility differential sites in female LOAD non-neuronal cells that overlapped LOAD-GWAS regions **(Table 3, Fig. S9)**. We focused the analysis on two representative LOAD differential chromatin accessibility sites in female LOAD non-neuronal cells that overlapped LOAD-GWAS loci: (1) LOAD more accessible sites surrounding the *ECHDC3* locus (annotated by the proximate gene to the associated SNP) and (2) LOAD less accessible sites surrounding the *ABCA7* locus. The loci were defined by anchoring on the associated SNPs +/- 500kb, rs7920721 (*ECHDC3*) and rs3752246 (*ABCA7*) using the UCSC Genome Browser^47^ (http://genome.ucsc.edu/) GRCh38/hg38 assembly released December 2013 **(Table 3, Fig. S9)**. Single-nuclei (sn)RNA-seq data was collected from female brains using a small subset group of two LOAD and two control (**Table S2,** marked with **\***), and differential expression analysis of female LOAD nuclei compared to control nuclei was performed for all genes mapped with these regions in three cell-type specific groups: 1) NeuN-non-neuronal, 2) Human Primary Cell Atlas (HPCA)-annotated astrocyte, and 3) HPCA-annotated microglia. Pseudogenes, RNA genes, and novel transcripts were excluded from the analysis. Out of 54 total genes within 1Mb of the two SNPs, 8 within *ECHDC3* locus and 46 within *ABCA7* locus (**Fig. S9**), many were up- or down-regulated as predicted by chromatin accessibility profiles (*i.e*. increased chromatin accessibility overlapping promoters and enhancers was associated with upregulation and vice versa). Overall 28 (51.9%), 23 (42.6%), and 24 (44.4%) genes of the NeuN-, astrocyte, and microglia clusters, respectively, showed differential expression with trends corresponding to the changes in chromatin accessibility. Of those genes, some showed nominal significance (6 (21.4%) for NeuN-, 3 (13.0%) for astrocyte, and 5 (20.8%) for microglia clusters), but did not reach adjusted statistical significance, while other trends did not reach nominal significance. This could be explained by the small sample size and/or number of cells in the clusters. Furthermore, we examined the consistency between our dataset and a similar recently reported dataset^28^. To address this question we conducted differential expression analysis for HPCA-annotated neurons, astrocytes, and microglia and compared the results for the top 15 significant genes, *i.e*. 5 from each cell type identified by the previous study^28^. We found that out of these 15 genes, our results show the same directionality for 13 genes, 10 of which also had significant adjusted p-values (**Table S14**), providing further validation of our findings.

**Table 3.** Differential snRNA-seq expression from level 3 female glia controls vs cases that are within 500 kb of LOAD-GWAS SNPs at loci found to be more accessible (rs7920721) or less accessible (rs3752246) in LOAD by ATAC-seq.

| SNP<br>(coordinates,<br>proximate gene) | Genes | NeuN- non-neuronal |  | HPCA-annotated astrocytes |  | HPCA-annotated microglia |  |
| --- | --- | --- | --- | --- | --- | --- | --- |
|  |  | Mean log(FC) | Adjusted *p* value | Mean log(FC) | Adjusted *p* value | Mean log(FC) | Adjusted *p* value |
| **rs7920721**<br>(chr10:11,678,309,<br>*ECHDC3*) | *CELF2* | <u>0.087</u> | 1 | -0.090 | 1 | <u>0.47</u> | **4.90 x 10<sup>-16</sup>** |
|  | *USP6NL* | <u>0.12</u> | 0.32 | -0.022 | 1 | <u>0.29</u> | 1 |
|  | *ECHDC3* | <u>0.067</u> | 1 | <u>0.016</u> | 1 | <u>0.31</u> | 1 |
|  | *UPF2* | -0.34 | **0.00042** | -0.31 | 1 | -0.48 | 1 |
|  | *DHTKD1* | -0.16 | 1 | -0.23 | 1 | <u>0.011</u> | 1 |
|  | *SEC61A2* | <u>0.028</u> | 1 | -0.028 | 1 | <u>0.15</u> | 1 |
|  | *NUDT5* | -0.032 | 1 | -0.097 | 1 | <u>0.21</u> | 1 |
| **rs3752246**<br>(chr19:1,056,493,<br>*ABCA7*) | *BSG* | 0.035 | 1 | 0.10 | 0.32 | <u>-0.11</u> | 1 |
|  | *POLRMT* | <u>-0.047</u> | 1 | <u>-0.030</u> | 1 | 0.061 | 1 |
|  | *FSTL3* | <u>-0.12</u> | 0.065 | <u>-0.13</u> | 1 | 0.047 | 1 |
|  | *PALM* | <u>-0.071</u> | 1 | 0.010 | 1 | <u>-0.25</u> | 1 |
|  | *PTBP1* | <u>-0.034</u> | 1 | <u>-0.053</u> | 1 | 0.065 | 1 |
|  | *AZU1* | <u>-0.012</u> | 1 | <u>-0.015</u> | 1 | <u>-0.039</u> | 1 |
|  | *MED16* | <u>-0.036</u> | 1 | <u>-0.051</u> | 1 | <u>-0.19</u> | 1 |
|  | *ARID3A* | 0.089 | 0.97 | 0.047 | 1 | 0.12 | 1 |
|  | *TMEM259* | 0.050 | 1 | 0.089 | 1 | <u>-0.11</u> | 1 |
|  | *ABCA7* | <u>-0.013</u> | 1 | <u>-0.022</u> | 1 | 0.075 | 1 |
|  | *STK11* | <u>-0.024</u> | 1 | <u>-0.041</u> | 1 | <u>-0.13</u> | 1 |
|  | *CBARP* | 0.013 | 1 | 0.017 | 1 | 0.026 | 1 |
|  | *ATP5F1D* | 0.040 | 1 | 0.11 | 1 | <u>-0.032</u> | 1 |
|  | *MIDN* | 0.042 | 1 | <u>-0.073</u> | 1 | 0.14 | 1 |
|  | *CIRBP* | <u>-0.71</u> | **1.52 x 10<sup>-57</sup>** | <u>-0.80</u> | **1.27 x 10<sup>-42</sup>** | <u>-0.51</u> | 1 |
|  | *C19orf24* | 0.033 | 1 | 0.014 | 1 | 0.12 | 1 |
|  | *NDUFS7* | 0.083 | **0.016** | 0.11 | **0.0014** | <u>-0.014</u> | 1 |
|  | *GAMT* | <u>-0.10</u> | 1 | <u>-0.065</u> | 1 | <u>-0.28</u> | 1 |
|  | *RPS15* | <u>-0.14</u> | 1 | <u>-0.19</u> | 0.66 | 0.088 | 1 |
|  | *APC2* | <u>-0.027</u> | 1 | 0.022 | 1 | <u>-0.14</u> | 1 |
|  | *PCSK4* | <u>-0.033</u> | 1 | <u>-0.025</u> | 1 | <u>-0.10</u> | 1 |

**Table 3** catalogues the genes surrounding the *ECHDC3* and *ABCA7* loci, excluding 26 genes that were not detected or had low fold change values. Out of seven genes positioned within the 1Mb region of the LOAD associated *ECHDC3* SNP, we found that *CELF2* was significantly upregulated in LOAD HPCA-annotated microglia clusters (mean log(fold change) = 0.47, adjusted *p* = 4.90×10^-16^, **Table 3**). In addition, out of 21 genes mapped within the 1Mb region of the LOAD associated *ABCA7* SNP, *CIRBP* was significantly downregulated in LOAD NeuN-non-neuronal clusters (mean log(fold change) = −0.71, adjusted *p* = 1.52×10^-57^, **Table 3**) and HPCA-annotated astrocyte clusters (mean log(fold change) = −0.80, adjusted *p* = 1.27×10^-42^, **Table 3**). These results demonstrated that alteration in chromatin accessibility in LOAD-GWAS loci correlates with changes in gene expression. Furthermore, the results suggested that the target genes within LOAD-GWAS loci affected by LOAD specific changes in chromatin state may not be simply interpreted as the most proximate gene to the associated SNP.

The intersection of LOAD- and glia-specific ATAC-seq sites and snRNA-seq patterns discovered novel LOAD regulatory elements that lead to gene expression changes in LOAD. Furthermore, intersecting these data sets provided a functional validation for the top significant findings from the ATAC-seq experiments showing sex-dependent LOAD-specific open and closed chromatin accessibility sites in non-neuronal cells. Collectively, these results suggest that integrating brain cell-type specific ‘omics data is a powerful mechanistic strategy to discover regulatory elements that impact expression of disease genes in a cell-type specific manner.

## Discussion

Decoding the genetic and genomic mechanisms of LOAD is a major challenge in the post-GWAS era, since the majority of the LOAD associated SNPs are in noncoding regions^1–7^. Noncoding disease-associated loci have been shown to be enriched for regulatory elements in tissues and cells relevant to the disease^37, 48–50^. Thus, post-GWAS research requires an in-depth characterization of cell-type specific DNA regulatory elements. Mapping chromatin accessibility has been widely used to identify the location of active DNA regulatory elements, including promoters, enhancers, and insulators^51–54^. We performed the first systematic interrogation of the chromatin accessibility landscape in neuronal and non-neuronal sorted nuclei from LOAD and healthy brains.

To our knowledge, this study represents the first and the most comprehensive dataset to date of chromatin accessibility in LOAD brains and matched controls. Furthermore, this study was performed with brain cell-type specific resolution, *i.e*. neuronal and non-neuronal cells. The study has four major findings for the field of LOAD epigenetics. First, we have generated a map of LOAD-associated cell-type specific chromatin accessibility sites. Second, we provide a catalogue of female-specific disease-associated chromatin accessibility sites in non-neuronal cells. Third, we suggest a non-coding regulatory mechanism, namely chromatin accessibility, by which ~25% of LOAD-GWAS loci may exert their effect. Fourth, we have demonstrated that LOAD associated changes in chromatin accessibility can result in gene dysregulation through their overlap with the transcriptome profile of LOAD-GWAS loci. Overall, these results suggest that the *cis* interactions between regulatory elements and key genes contribute, at least in part, to the development and/or progression of LOAD. Of note, 87% of the differentially accessible chromatin peaks in the female NeuN- analysis and 92% of the differentially accessible chromatin peaks in the NeuN+ analysis fall within known regions of topologically associating domains (TADs) from fetal brain^55^. This suggests that chromatin regions associated with risk for LOAD belong to highly interacting regulatory domains. Furthermore, several LOAD loci may exert their pathogenic effects in a cell-type specific manner while others act in multiple cell types to drive LOAD pathogenesis. Alternatively, these cell-type specific and common changes in chromatin structure may represent secondary effects as consequences of disease-related processes such as neurodegeneration and gliosis.

A growing number of epigenome-wide association studies (EWAS) in LOAD have profiled DNA methylation, hydroxymethylation, and histone acetylation marks (H4K16ac), and assessed associations with LOAD risk and other related endophenotypes including the burden of pathology^17–24^. These studies have been a powerful approach to validate known LOAD loci, discover new candidate genes, and identify disease-related pathways^17–24^. However, the majority of the LOAD EWAS datasets were generated using bulk brain tissues, and the heterogeneous cell-type populations of brain tissue samples pose technical and biological limitations. Our results showed that a substantially larger number of ATAC-seq sites are uniquely detected in the neuronal and non-neuronal cells, compared to bulk tissues obtained from the same donors. This observation demonstrates the importance of cell-type specific epigenomic studies relative to bulk tissues, as it reduces cellular heterogeneity allowing the identification of more cell-type specific signals.

Collectively, our outcomes open a new window for the exploration of the particular cell-types that contribute to LOAD pathogenesis, and the genes and pathways that mediate the cell-type specific pathogenic effects.

These data advance the mechanistic understanding of LOAD, and moreover, uncover new candidate LOAD loci. To date, in addition to this study only five others (two transcriptome, two DNA-methylation, and one histone acetylation) have compared genomic signatures stratified by different cell types in the LOAD brain using sorted- and single-nuclei based methodologies^27 28 29 51 56^. Altogether, our and other studies demonstrated the importance of applying these cell-type specific approaches in molecular analyses of brain tissues and highlight the impact of transitioning into single-cell based ‘omics studies in LOAD functional genomic research. In addition, we show that integrating our cell-type specific ATAC-seq data with our scRNA-seq LOAD data corroborate the interpretation of the results and provide functional validation. By aligning the two datasets, we identified LOAD-specific female-dependent non-neuronal regulatory elements around two differentially expressed genes in LOAD glia cells. Moreover, analysis of chromatin accessibility differential sites in female LOAD non-neuronal cells that overlapped LOAD-GWAS regions not only provided functional validation and established the link between LOAD -GWAS -ATAC-seq and -snRNA-seq but also demonstrated that genes other than the most proximate to the associated SNP may play a role in LOAD pathogenesis. These results exemplify the potential of integration of cell-type specific datasets to validate known LOAD loci and also to identify new candidate genes. In future studies, a larger sample size may allow conducting a chromatin accessibility QTL study to determine colocalization with GWAS loci.

Several bulk tissue ChIP-seq studies have used functional genomics and integrative systems biology approaches to infer cell types. Consistent with our findings, these epigenomic studies strongly suggest that non-neuronal cell types contribute to LOAD-specific histone marks associated with active regulatory elements (promoters and enhancers). It was reported that LOAD GWAS loci were enriched in enhancer elements specific to immune cells^57^ and tangle-associated H3K9ac signals located in both promoters and enhancers were significantly associated with modules classified as non-neuronal^58^. Recently, two studies used FANS-sorted nuclei followed by ChIP-seq and demonstrated that microglia were the non-neuronal cell type contributing to LOAD epigenomic signatures. Nott *et al*. found that LOAD SNP heritability was most significant in microglial enhancers^59^, and Ramamurthy *et al*. showed that hyperacetylated peaks in microglia colocalize more with LOAD SNPs than the histone acetylome of other cell types^56^. Collectively, ours and others’ studies point to non-neuronal epigenomic dysregulation, likely microglial, as a major contributing factor to LOAD pathogenesis.

Sex is an important factor in LOAD etiology and there are sex differences in disease risk, progression and clinicopathological phenotypes^60^. Foremost, there is a sex-dependent difference in LOAD prevalence and almost two-thirds of LOAD patients are female^61^. Historically, it has been attributed to the longer average life expectancy in women^62^, however, recent research suggests that other factors, such as the sudden drop in the level of sex hormones (estrogens) in women at menopause, contribute to the differences in susceptibility between males and females^42^. It was also reported that women manifest faster disease progression and cognitive decline, increased brain atrophy and pathological burden largely driven by neurofibrillary tangles^42^, and a more advanced disease stage as indicated by CSF biomarkers, especially higher concentrations of total tau and phosphorylated tau. Although several conflicting reports suggested opposite trends^63^, the effect of sex on LOAD has been widely accepted. Nonetheless, the molecular mechanisms underlying the role of sex as a risk factor in LOAD are understudied. Our study provides new insights into these gaps in knowledge showing sex-dependent changes in chromatin structure between LOAD and control brains. We identified hundreds of LOAD differential chromatin accessibility sites specific to females, which overlap nearly one-third of all LOAD GWAS regions. Since the majority of differential chromatin accessibility sites do not overlap LOAD-GWAS regions, these represent novel candidate LOAD loci. Moreover, a female-specific effect on LOAD-associated changes in chromatin accessibility appeared exclusively in glial cells and resulted in nearly three-fold overrepresentation of sites that were more closed in female LOAD patient samples. However, we cannot rule out the possibility that LOAD-associated changes in chromatin accessibility also occur in glia from male LOAD patients, but because of the plausible much smaller effect size in males, we could be underpowered to detect significant associations in our male cohort. These results warrant further investigation to determine the effects of sex-dependent chromatin remodeling on dysregulation of gene expression in the context of LOAD. In this respect, the recent scRNA-seq study^28^ also reported sex-dependent LOAD effects and specifically observed a sex-specific differential transcriptional response to LOAD pathology, enrichment of females cells in LOAD-associated cell subpopulation, and higher expression in females for the marker genes of LOAD-associated cellular subpopulations^28^. Future integration of diverse ‘omics datasets stratified by sex will decipher the underpinning mechanisms of sex differences in LOAD by establishing the cross interactions between sex-dependent chromatin structure and function, transcriptome and LOAD phenotypic outcomes.

One of the loci that showed sex-dependent effects in our study is the *APOE* linkage disequilibrium (LD) region. The e4 allele of the *APOE* gene is the first identified, most highly replicated, and the strongest genetic risk factor for LOAD^64, 65^. Furthermore, LOAD GWAS have confirmed strong associations with the *APOE* LD genomic region, and no other LOAD-association remotely approached the same level of significance ^3, 66, 67^ Interestingly, female carriers of *APOE* e4 have an increased risk of LOAD versus male and the adverse effect of *APOE* e4 on LOAD biomarkers was generally stronger in women versus men^68–72^. Overall the *APOE* LD region displayed more open chromatin in glia versus neurons as expected since this gene is much more highly expressed in glia. Interestingly, we found decreased chromatin accessibility at multiple sites in female LOAD glia across the *APOE* region. This result provides molecular clues to the observations that *APOE* e4 allele confers a greater risk for LOAD in women than in men^68–72^. In addition, significant downregulation of *APOE* in astrocytes from LOAD brains, although not sex-dependent, was reported by scRNA-seq analysis. This evidence is consistent with the trend we found of more closed chromatin in LOAD samples, providing functional validation to our result^28^. In summary, we proposed molecular insights based on chromatin structure that may explain, at least in part, some aspects related to the role of *APOE* in LOAD.

In conclusion, this LOAD genomic research pioneers the approach of brain cell-type specific chromatin accessibility profiling and lays the foundation for additional sorted- and single-nuclei ‘omics analyses in LOAD. Our outcomes warrant further investigations using a larger sample size to enhance the discovery smaller LOAD-associated effects on chromatin accessibility and to allow the utilization of a single peakset and set of covariates to perform multiple testing. Future and ongoing studies using even more advanced single-cell technologies will generate complementary ‘omics datasets with finer cell-type resolution from larger well-characterized LOAD cohorts. Data sharing via publicly available portals, such as Accelerating Medicines Partnership-Alzheimer’s Disease (AMP-AD), will facilitate integrative single-cells ‘omics towards moving forward our understanding of the underpinning genetic drivers and molecular mechanisms of LOAD.

## Supplemental Data

Supplemental Data include 12 figures and 15 tables.

## Declaration of Interests

The authors declare no competing interests.

## Acknowledgments

We thank the Kathleen Price Bryan Brain Bank at Duke University (funded by NIA AG028377) for providing us with the brain tissues, the Duke Cancer Institute Flow Cytometry Shared Resource for sorting, and the Duke Sequencing and Genomic Technologies Shared Resource for sequencing. We also thank John Ervin for his assistance in obtaining the brain samples with neuropathological and clinical data required for the study, and Lynn Martinek for providing training and technical assistance in using the cell sorter and in analyzing the flow data. This work used a high-performance computing facility partially supported by grant 2016-IDG-1013 (“HARDAC+: Reproducible HPC for Next-generation Genomics”) from the North Carolina Biotechnology Center. The results published here are in part based on data obtained from the AD Knowledge Portal (https://adknowledgeportal.synapse.org). Study data were provided by the Rush Alzheimer’s Disease Center, Rush University Medical Center, Chicago. Data collection was supported through funding by NIA grants P30AG10161, R01AG15819, R01AG17917, R01AG36836, and the Illinois Department of Public Health. Additional phenotypic data can be requested at www.radc.rush.edu.

## Funding

This work was funded in part by the National Institutes of Health/National Institute on Aging (NIH/NIA) [R01 AG057522 to O.C-F.] and a seed grant from The Duke Center for Genomic and Computational Biology (to O.C-F. and GEC).

## Author contributions

G.E.C. and O.C-F. conceived of the presented idea. J.B. and O.C-F. acquired brain samples, designed the study sample, and interpreted metadata. J.B. and Y.Y. performed brain tissue handling, nuclei extraction, labeling and sorting, and fluorescence microscopy evaluation to ensure nuclei were of high quality. I.P., D.S., and D.C. provided technical support in extraction of bulk nuclei from brain tissues. A.S. performed the ATAC-seq experiments. J.B., Y.Y. and J.G. performed the snRNA-seq experiments. M.E.G. and A.A.K. performed QC and covariate analysis. L.S. performed alignment, peak quantification, QC assessment, differential chromatin, and motif analysis. R.G. advised on the motif analysis. J.G. and H.F. performed the snRNA-seq data analysis and differential expression analysis. J.L. assisted with the QC metric analysis. K.S. assisted with whole genome genotyping. A.A.K., G.E.C. and O.C-F. planned and supervised the work. G.E.C. and O.C-F. obtained funding support. All authors discussed and interpreted the results. J.B., L.S., A.S., Y.Y., M.E.G., J.G., A.A.K., G.E.C., and O.C-F. designed the study and wrote the manuscript. All authors read and approved the manuscript.

## Data Availabillity

The ATAC-Sequencing and single-nucleus RNA-Sequencing data are available at Synapse (https://www.synapse.org/#!Synapse:syn20755767). The DOI for this dataset is 10.7303/syn20755767. The data are available under controlled use conditions.

## Notes

### Competing Interest Statement

The authors have declared no competing interest.

