## Supplemental Figures S1-S12 for "Sex dependent glial-specific changes in the chromatin accessibility landscape in late-onset Alzheimer’s disease brains"

**Figure S1**: Pearson correlations of all potential numerical covariates and the first ten principal components of the potential numerical covariates.

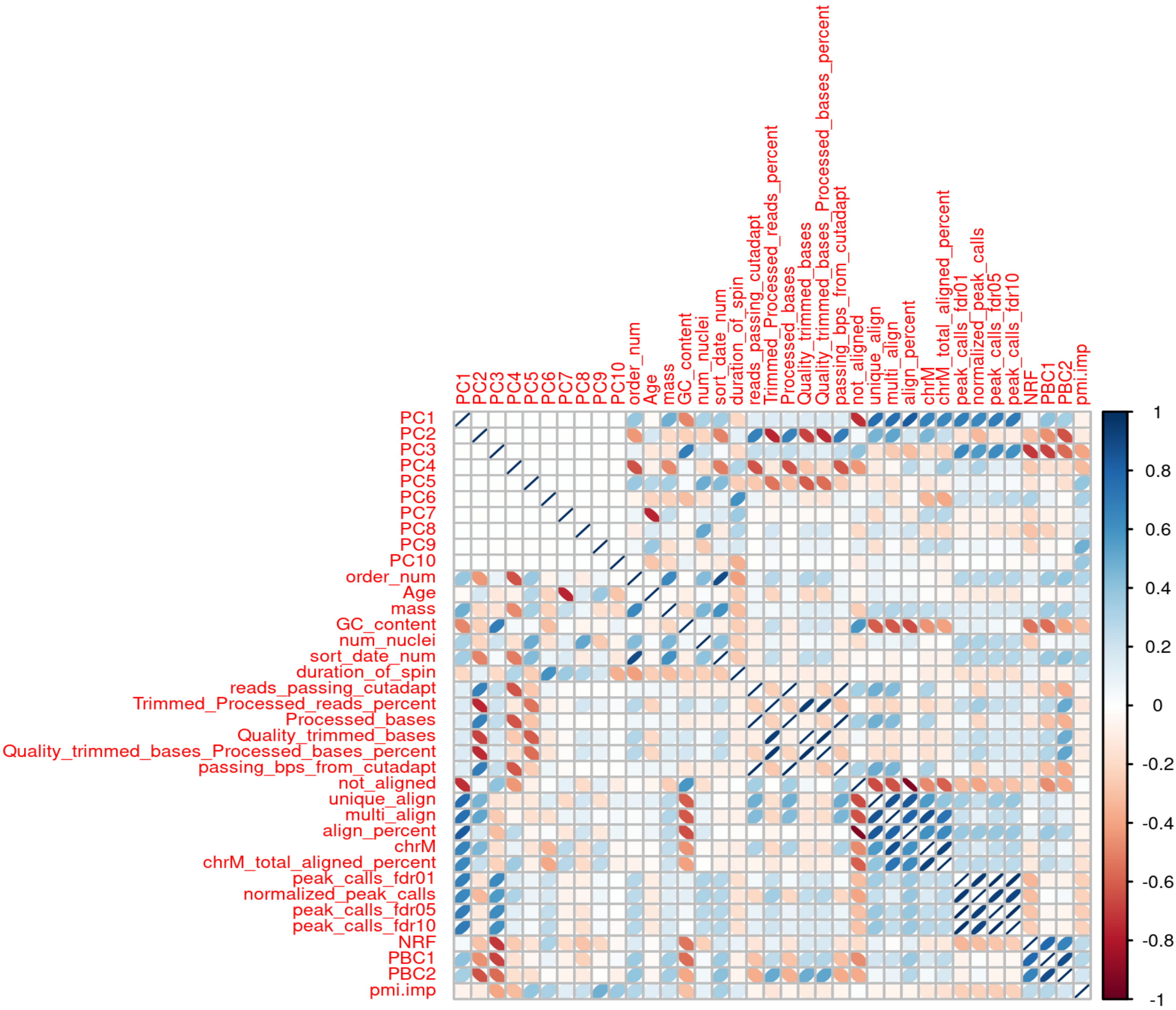

**Figure S2**: Association of meta variables with case-control status at a Bonferroni significance level of q<0.05. The vertical line indicates Bonferroni significance. Linear regression was performed for numerical variables (n=27) and chi-square tests were conducted for categorical variables (n=2, excluding diagnosis).

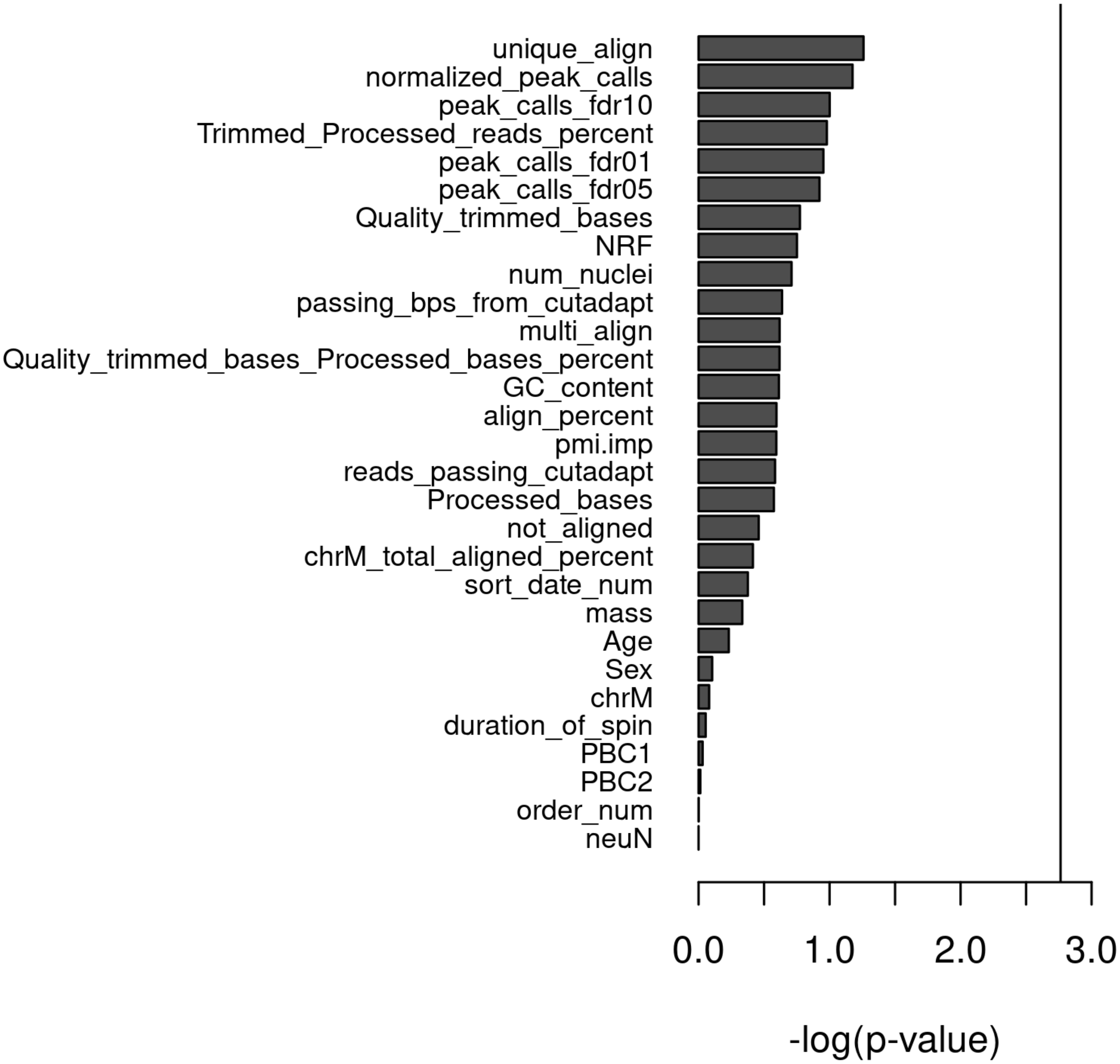

**Figure S3:** Scree plot depicting the proportion of total variance explained by each principal component of peaks in all samples passing QC.

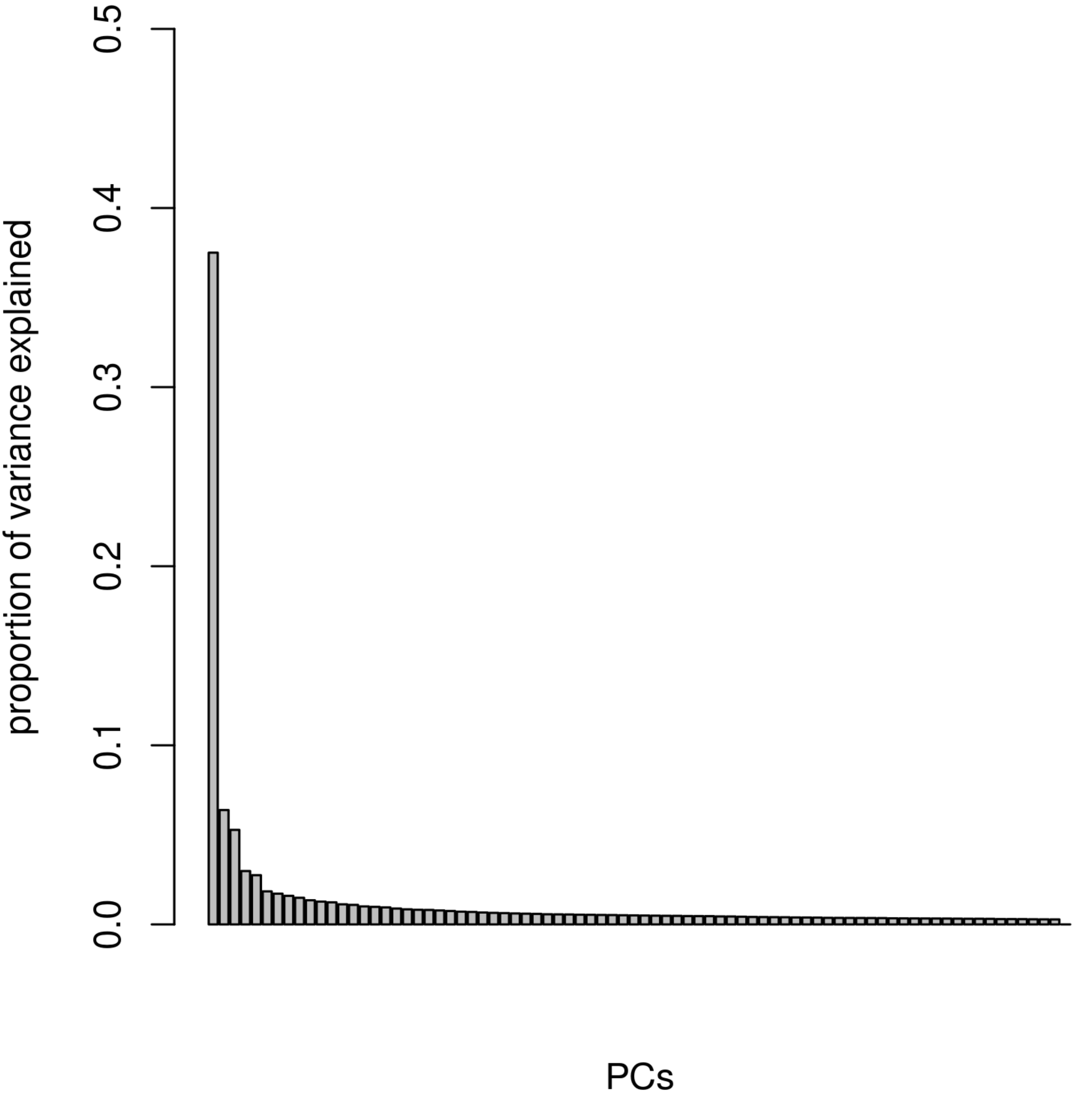

**Figure S4**: Level 1 comparison: Genomic distribution of Top 5000 differential sites of neuron/non-neuron.

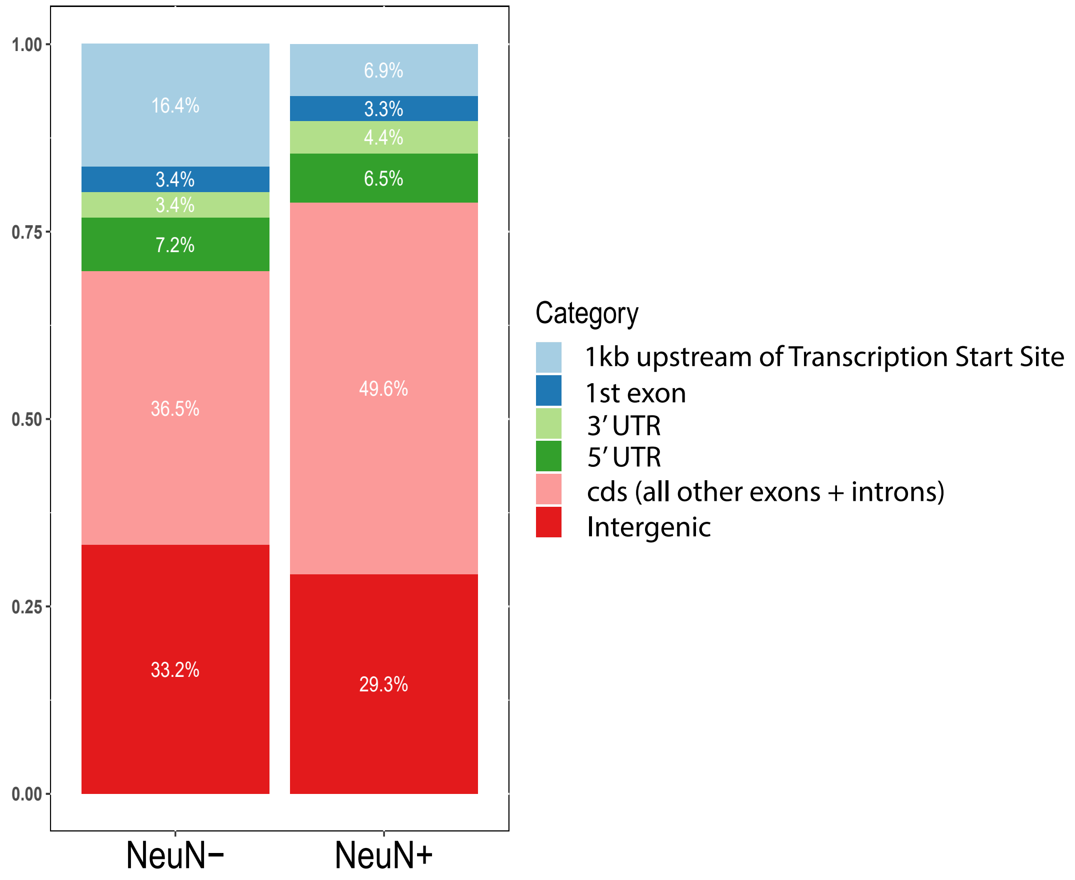

**Figure S5:** Fullard et al. vs. control samples from this study. (**a**) Venn diagram of differential sites from Fullard et al. publication and this study. Differential sites were generated by DESeq2 at fdr 0.01. Upper panels were analyzed by choosing the same numbers of samples from this study as those in Fullard et al. publication. (**b**) Peak length distribution of Fullard et al. 8+8, This study 8+8 and This study 26+21

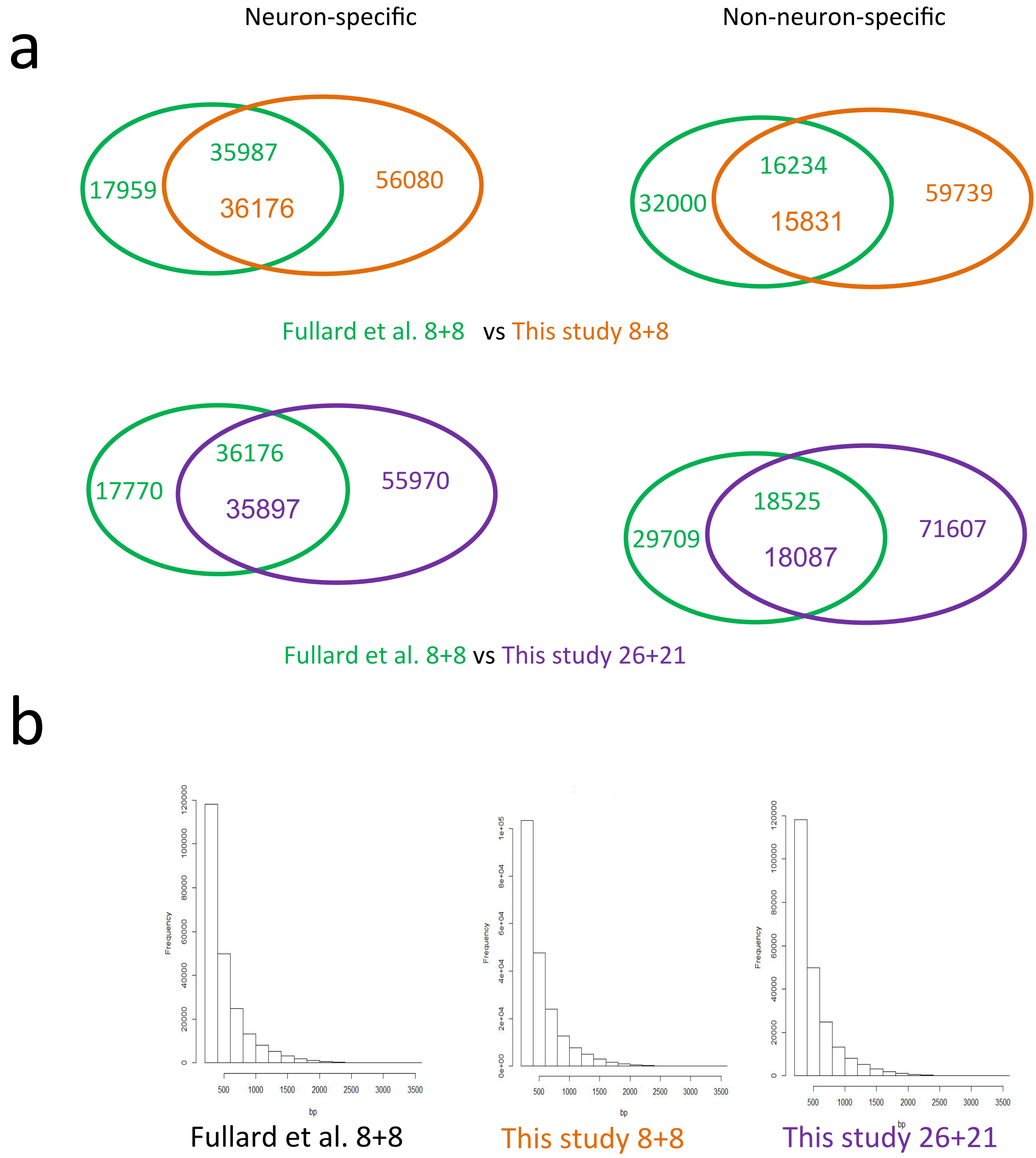

**Figure S6:** MA plots (FDR < 0.05) of Diff analysis of Whole tissue vs neuron/non-neuron mixed (n= 6 vs 6). (**a**) Comparison of sorted neuron vs whole tissue. (**b**) Comparison of non-neuron vs whole brain tissue. (**c**) Comparison of 1:1 in silico mixed neuron/non-neuron cells control vs cases (no differential sites, dispersion= 0.23). (**d**) Overlap between DE sites

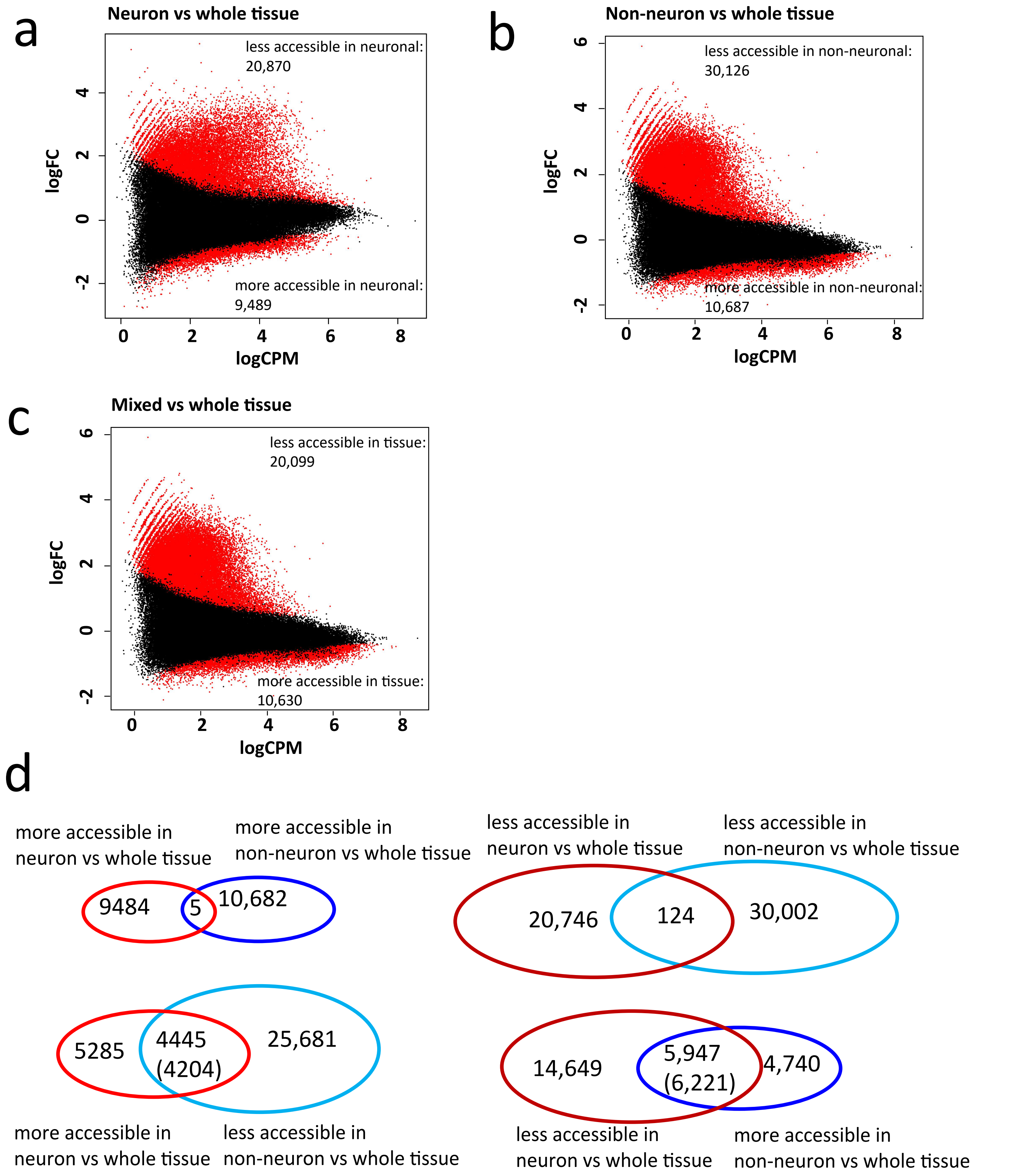

Figure S7: Box plots showing accessibility of peaks not differential between cases and controls for (a) CLU, PTK2B, (b) APOE, and (c) IQCK. The green transparent box indicates the regions for the controls in order. Box plots show the median, 25th percentile, and 75th percentile. Box plot whiskers show the 75th and 25th percentiles plus and minus 1.5 times the interquartile range, respectively.

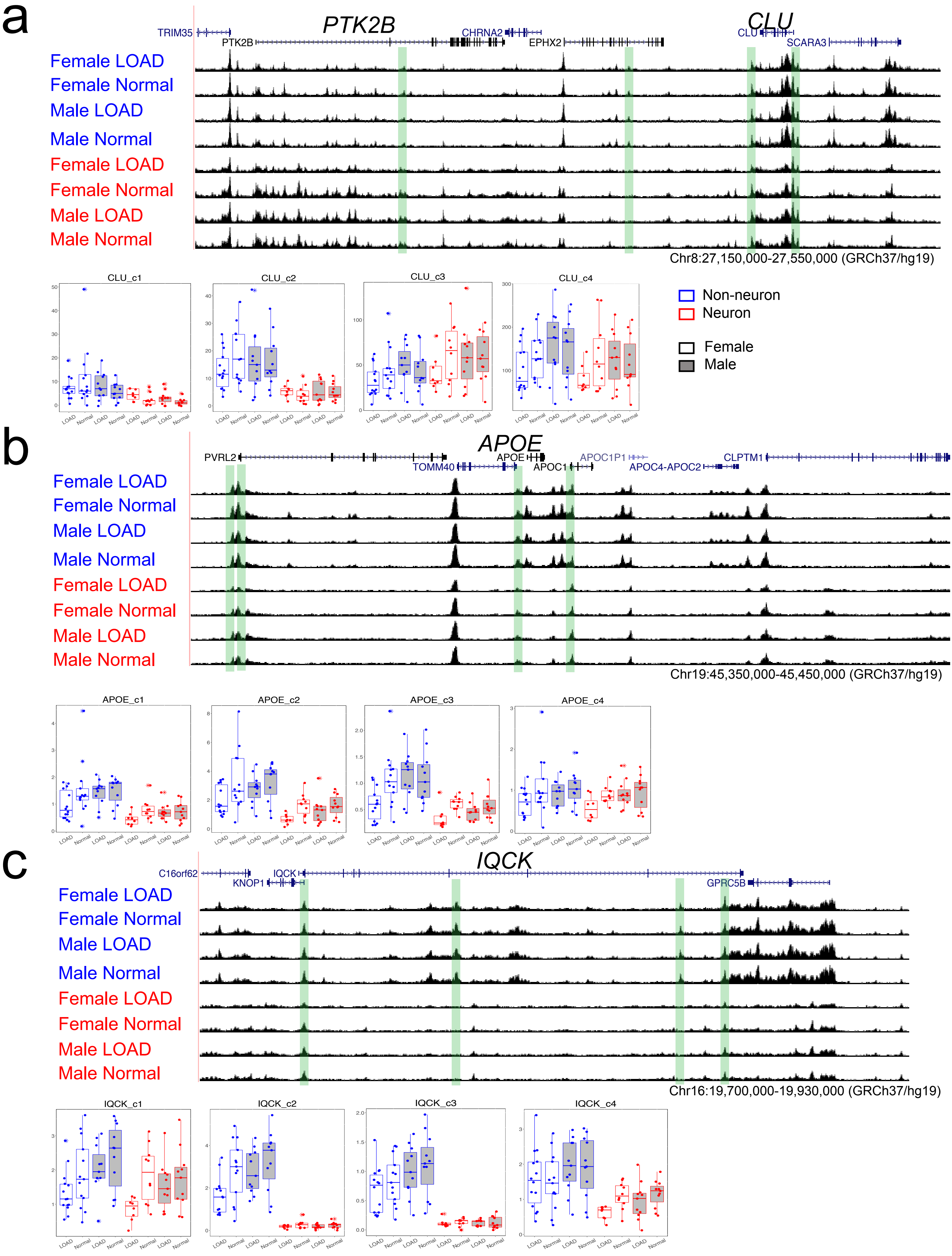

**Figure S8:** Permutation (n=10000) controls for differential sites overlap with GWA LOAD sites. Red lines represent LOAD sites numbers overlapped by differential sites. **a.** neuron LOAD up, randomly chosen 537 peak calls (pvalue=0.13); **b.** neuron LOAD down, randomly chosen 947 sites(pvalue = 0.06); **c.** female non-neuron LOAD up , randomly chosen 1000 sites (pva= 0.98); **d.** female non-neuron LOAD down , randomly chosen 2352 sites (pva= 0.05);

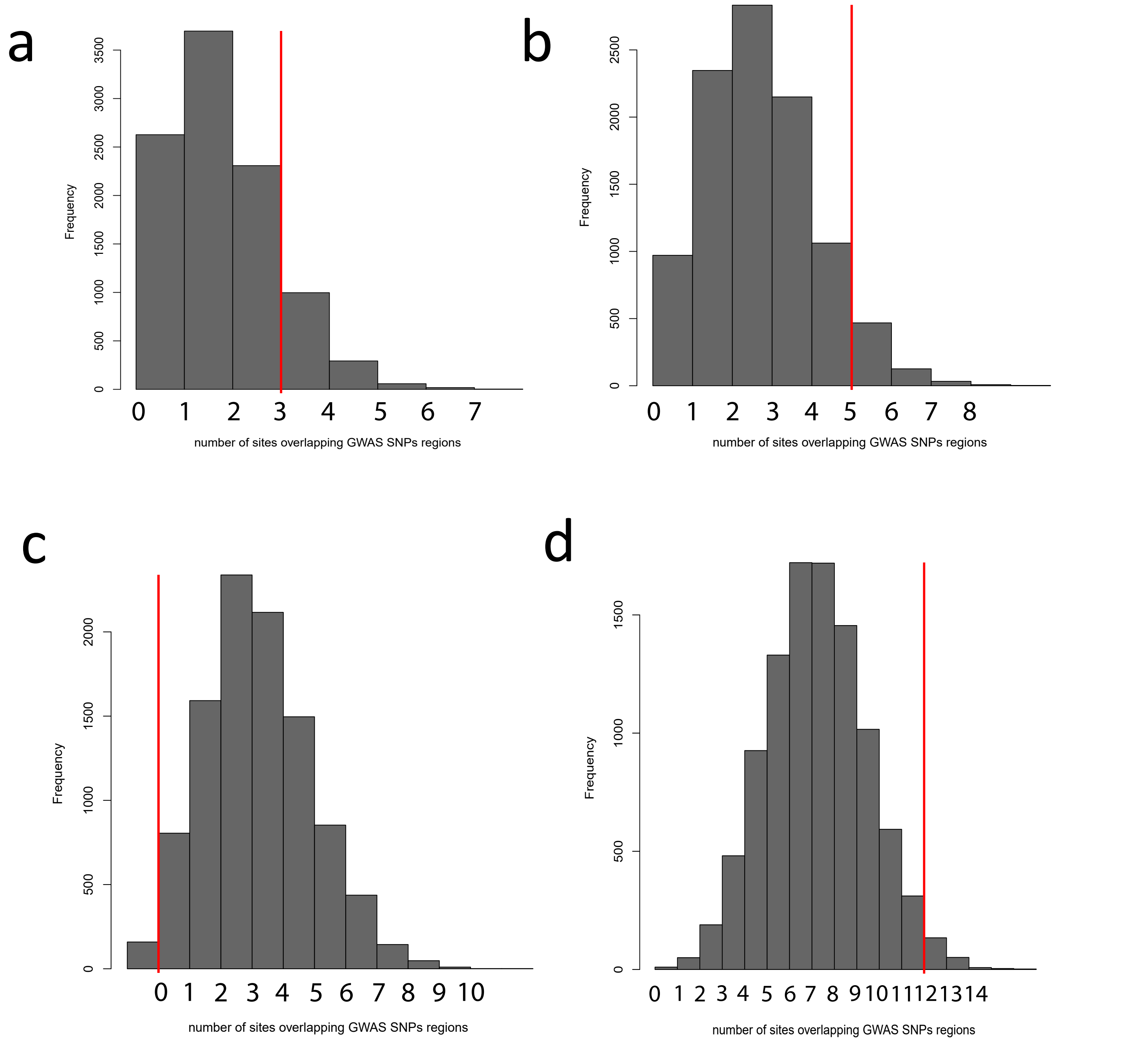

**Figure S9:** All genes within 1Mb of LOAD-GWAS loci found to be more (top) or less (bottom) accessible by ATAC-seq of FANS-sorted nuclei. SNPs (light blue lines) were anchored in the center of the region and red boxes indicate genes found to be significantly dysregulated by snRNA-seq. Pseudogenes, RNA genes, and novel transcripts are excluded. Figures generated using the UCSC Genome Browser (http://genome.ucsc.edu/) GRCh38/hg38 assembly released December 2013.

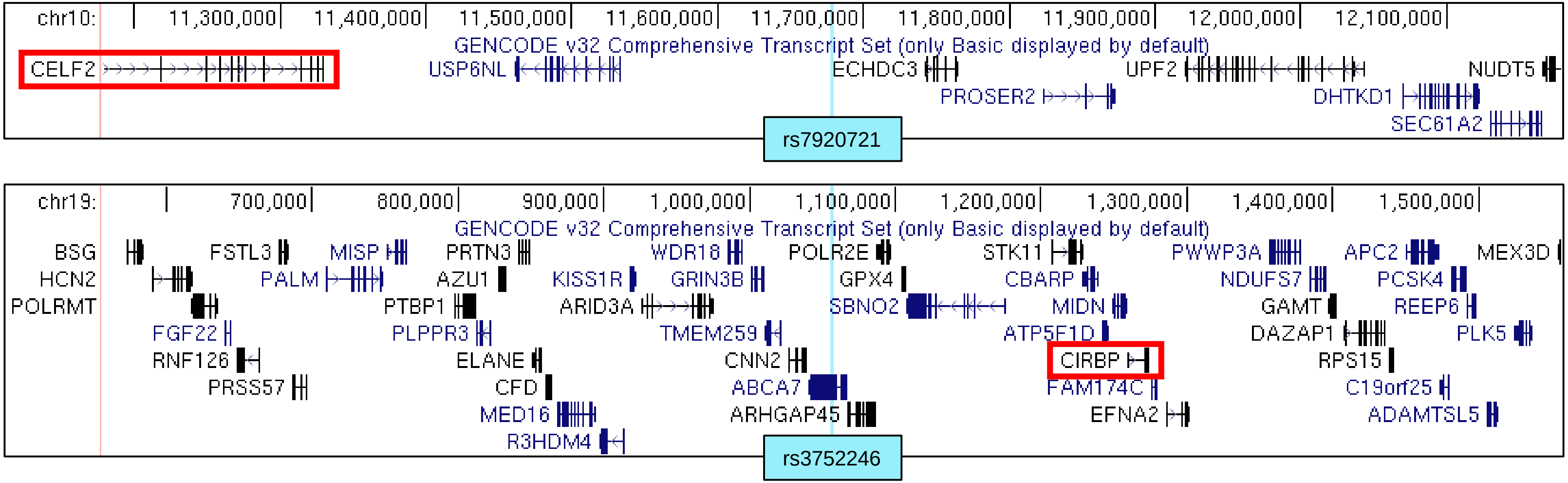

**Figure S10:** Clustering of 90 replicates for data selection. (**a**) Hierarchical clustering. (**b**) K-mean clustering(K=2).

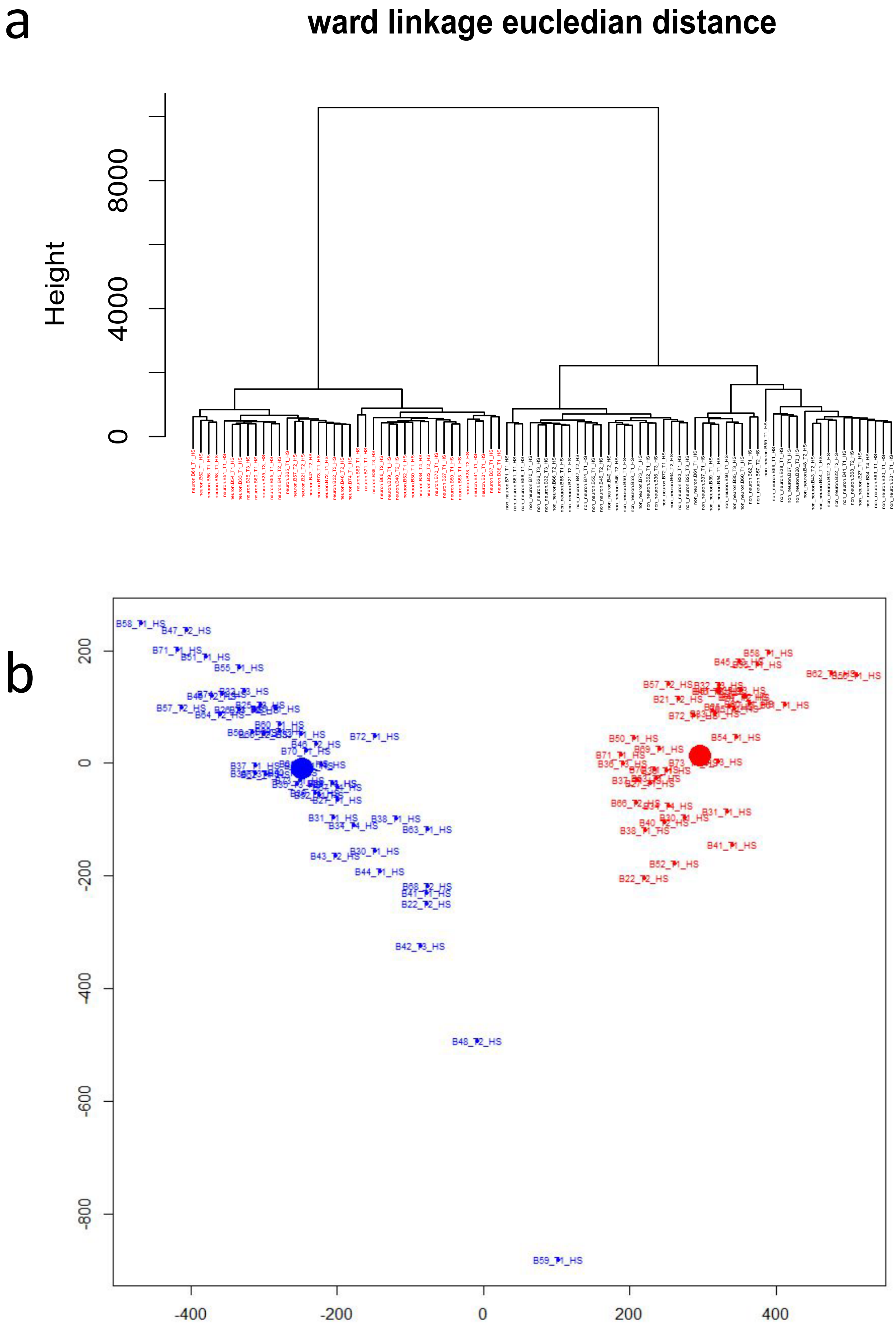

**Figure S11**: Scatterplot of PC1 vs. PC2 peaks showing association between PC1 and cell type ($p=5.11 \times 10^{-45}$).

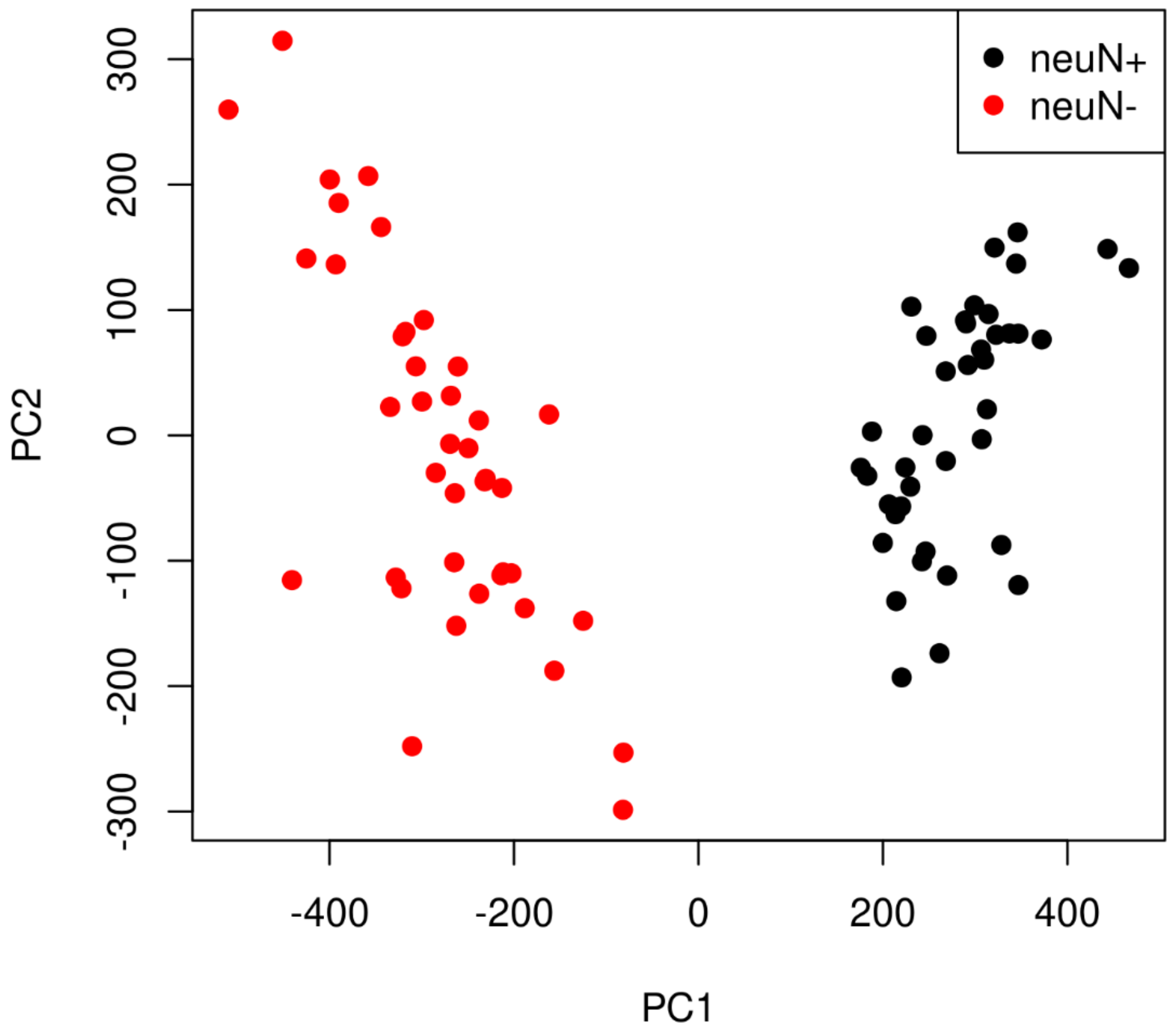

**Figure S12**: (**a**) Differential sites found in the female glia normal vs LOAD comparison (blue and red circles) were mapped to MA plots of female neuron normal vs LOAD comparison (black dots). For the female glia normal vs LOAD comparison, blue circles represent differential sites which were more accessible in LOAD while red circles represent differential sites which were less accessible in LOAD. (**b**) Differential sites found in 27 replicates of female glia normal vs LOAD comparison were mapped to log fold changes plot of 18 replicates of female glia normal vs LOAD comparison vs. 22 replicates of male glia normal vs LOAD comparison. Black dots represent sites shared by the above two comparisons. Blue circles represent differential sites which were corresponding to those in female glia normal vs LOAD and more open in LOAD. Red circles represent differential sites which were corresponding to those in female glia normal vs LOAD and less open in LOAD. The dashed line represents y=x. The solid line represents linear regression of all of data points.

**a** **b**

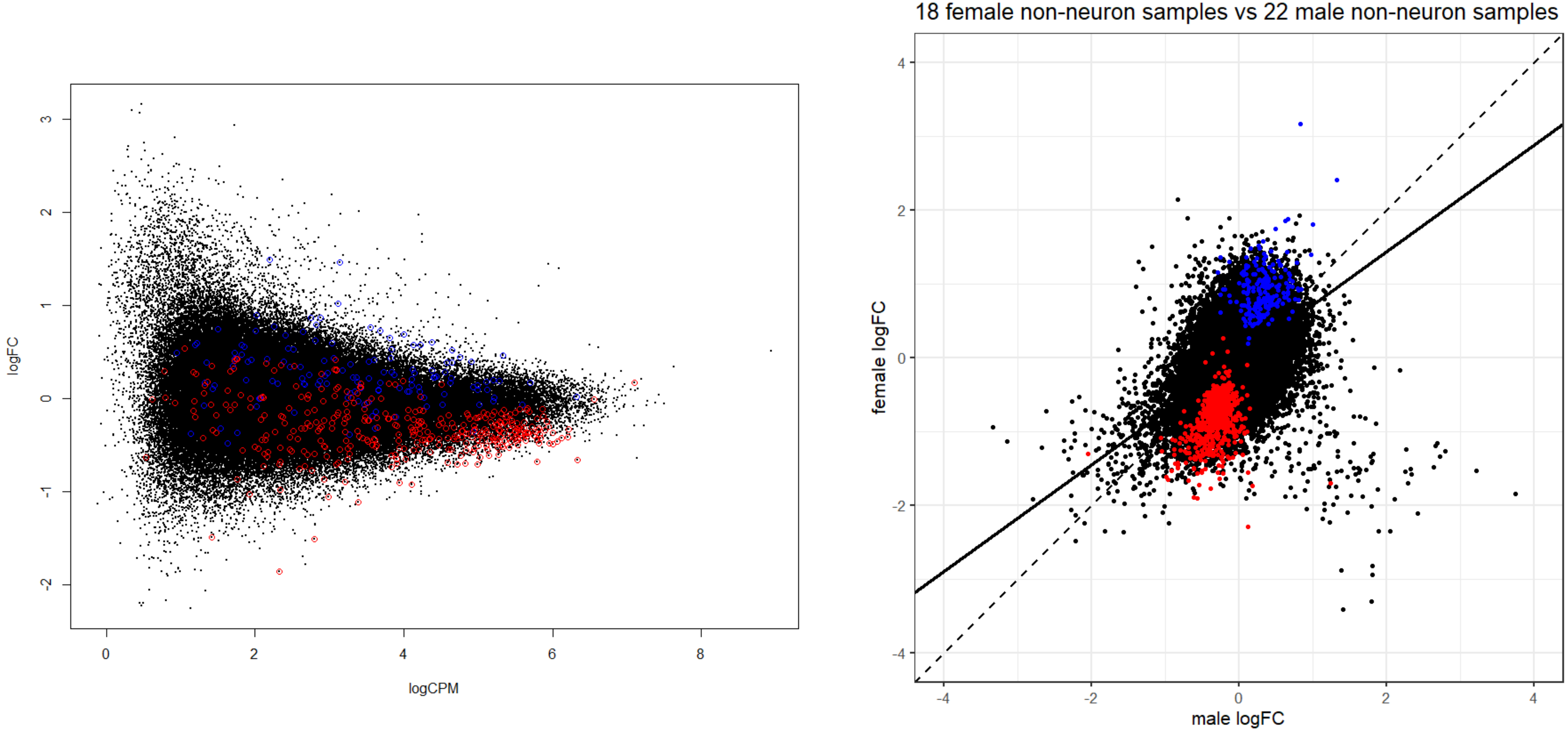
